# Diversity and spatiotemporal activity patterns of medium and large mammals in the Niokolo-Koba National Park, Senegal

**DOI:** 10.64898/2026.03.03.709351

**Authors:** Lisa Ohrndorf, Augustin Brouillet, Annika M. Zuleger, Ndiouga Diakhaté, Djibril Coly, Chérif Younousse Kéba Camara, Amadou Bamba Diedhiou, Irene Gutiérrez Díez, Julia Fischer, Dietmar Zinner

**Author notes:** Corresponding author: Lisa Ohrndorf.

## Abstract

West African savannahs provide habitats to diverse species assemblages, yet remain understudied compared to their East and Southern African counterparts. The Niokolo-Koba National Park in southeastern Senegal constitutes one of the largest remaining protected areas in West Africa and supports a mosaic of savannah and forest habitats with a diverse assemblage of medium- and large-sized mammals. Here, we analysed camera-trap data originally collected to monitor predator presence in the northwestern sector of the National Park. We deployed 37 cameras across 37 km² from February 2022 to March 2023, resulting in 13,161 camera-trap-days. We assessed alpha diversity indices and spatiotemporal activity patterns of large and medium-sized mammals across habitat types. Evenness values – the degree to which species abundances are distributed uniformly within a community – were higher in the savannah than in forest habitats, although overall species richness was comparable. In contrast, animal sighting rates were higher in forests than in savannahs. Estimated diel activity mostly corresponded with established species-specific behavioural patterns. Our analyses revealed differential use of certain habitat types across the day, likely driven by spatially segregated sleeping sites and foraging locations. Our results provide a reference for future studies and monitoring efforts and highlight the value of the forest-savannah mosaic for the local species assemblage within the larger ecosystem of Niokolo-Koba National Park.

## Introduction

Savannahs and savannah-forest mosaics are among the world’s largest biomes, covering up to 30% of the overall land surface (Shorrocks 2007, Bond 2019). Most of these savannahs are located in Africa, providing habitats for a wide range of medium and large mammal species. However, savannah ecosystems, especially in eastern and western Africa, are under increasing pressure, resulting in a sharp decline in large-mammal populations over recent decades and contributing to the global decline in biodiversity (Craigie et al. 2010). Habitat loss due to overexploitation and fragmentation is among the main drivers of this decline and disproportionately affects wide-ranging and large-bodied species (Scholte et al. 2025). Savannahs are thus central to conservation planning across Africa. At the same time, savannah ecosystems are often overlooked or take second place in conservation and restoration planning (Buisson et al. 2019, Dudley et al. 2020). While savannah ecosystems in East and Southern Africa have been extensively studied within protected areas, similar research in West Africa remains limited (Bauer et al. 2021). This lack of data is concerning, given the continued decline of mammal populations in the region and the growing need to identify and protect remaining areas of ecological importance (Craigie et al. 2010, Scholte et al. 2025).

The Niokolo-Koba National Park (NKNP), located in southeastern Senegal, is one of the largest remaining protected areas in West Africa (UNESCO World Heritage Centre 2025). It has been listed as a UNESCO World Heritage Site since 1981 due to its ecological importance (UNESCO World Heritage Committee 1981). The park contains a mosaic of savannahs, gallery forests and wetlands, and historically supports a wide range of mammal species, including the giant eland (*Taurotragus derbianus*), chimpanzees (*Pan troglodytes verus*), lions (*Panthera leo leo*), leopards (*Panthera pardus*), and elephants (*Loxodonta* sp., likely *L. cyclotis* although the taxonomy is uncertain, due to possible hybridisation between *L. africanus* and *L. cyclotis* (Dupuy 1971, Kuhner et al. 2025)). Over the past decades, NKNP has experienced a range of pressures, including poaching, livestock grazing, and the planned construction of a dam on the Gambia and Niokolo Rivers upstream of the park, which have severely threatened its water regime. In 2007, these challenges led to its inclusion on the UNESCO List of World Heritage in Danger (UNESCO World Heritage Committee 2007). Following increased conservation and monitoring efforts, the park was removed from the list in 2024 (UNESCO World Heritage Committee 2024, Houéhounha et al. 2025). Nonetheless, a range of threats to the park’s biodiversity persist, including the progressive drying of previously perennial water bodies due to invasive species such as *Mimosa pigra*, bush encroachment, illegal logging and grazing, mining, and associated water pollution (UNESCO World Heritage Committee 2024). These changes may have serious implications for wildlife, particularly for the park’s water regime during the dry season, when access to water becomes a limiting factor (Houéhounha et al. 2025, UNESCO World Heritage Committee 2025).

In this study, we analysed camera-trap data collected to monitor predator presence near Simenti, a ranger post in the northwestern sector of NKNP (Ohrndorf et al. 2025a, b). We explored these data to assess species diversity and spatiotemporal activity patterns of large and medium-sized mammals across habitat types. Specifically, we assessed which species were present and how frequently they were captured by the cameras year-round. We determined alpha diversity indices and local species detection rates to examine whether and how species detections varied across locations. Further, we investigated potential seasonal variation in habitat use across species. We then estimated temporal activity patterns of frequently detected species across the entire study site and across habitat types to gain insights into the spatiotemporal structure of habitat use of the local mammal community. We aim to provide high-resolution, up-to-date information on mammal diversity and activity patterns in this section of NKNP, supporting ongoing conservation efforts and contributing to a broader understanding of mammal ecology in the local habitat mosaic.

## Material and Methods

### Study site

Data collection for this study took place at the Centre de Recherche de Primatologie (CRP) long-term field site in Simenti (Fischer et al. 2017). The field site is located in southeast Senegal within the NKNP next to the Gambia River (Figure 1). The site lies within the Sudanian and Sahelo-Sudanian climatic zones and is characterised by pronounced seasonality and considerable rainfall variability (Arbonnier 2002). Average annual precipitation in Simenti is approximately 950 mm, with the rainy season typically lasting from June to October. May and mid-October are transitional periods with little rainfall (Arbonnier 2002). The vegetation is classified as a mosaic of grasslands, wooded savannahs, and gallery forests along streams and other perennial water bodies (Arbonnier 2002, Burgess et al. 2004).

**Figure 1:**
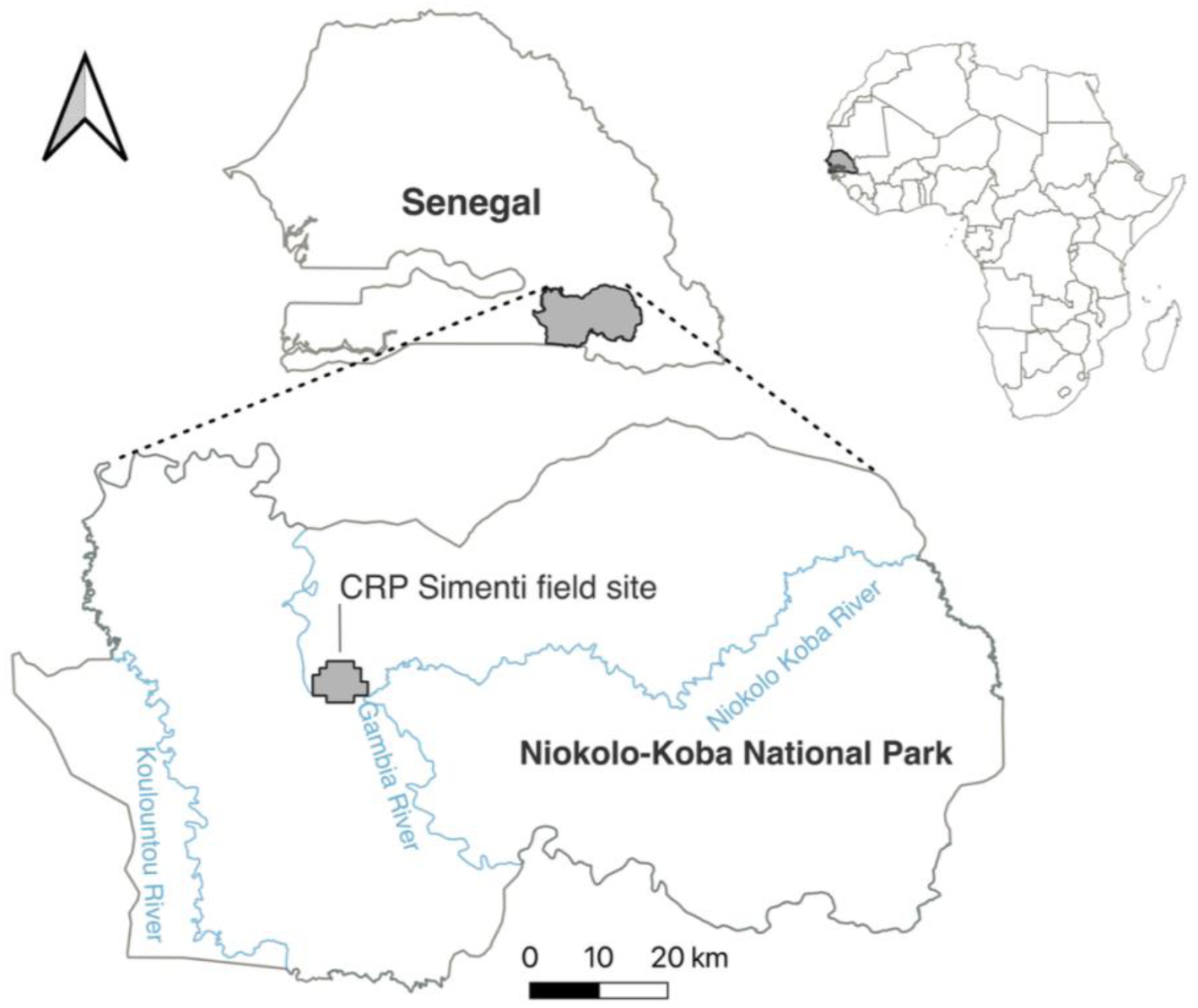
Location of the study site within the Niokolo-Koba National Park in southeast Senegal.

### Camera-trap setup

The data analysed in this study were collected using unbaited, motion-triggered camera-traps deployed from February 2022 to March 2023. To ensure systematic sampling across a wide area, the cameras were positioned in a grid with approximately one camera per km², resulting in 37 sampling points. In total, 45 camera-traps were available (33 SECACAM Pro Plus, 1 Braun Scouting Cam Black 1300, 3 TEC.BEAN SG-009, 3 Coolife Trail Camera, and 5 DÖRR Snap Shot Limited 5.0S). Of these, 37 were deployed simultaneously across sampling points, while the remaining eight served as backup in case of malfunction. All cameras, except one, were mounted on trees at approximately 0.5 to 0.6 m above ground level, facing visible animal trails. One camera, located on an open laterite airfield, could not be placed near an evident animal trail. The true camera-trap locations deviated on average by 16.9 m from the planned grid points, with only two exceeding a deviation of 50 m (Figure 3). To ensure continuous data collection, we replaced the batteries and memory cards of all cameras monthly. During the rainy season, we cleared a five-meter radius around each camera of grass and weeds to maintain unobstructed visibility and prevent fire damage. All cameras were programmed to take three consecutive pictures per trigger event, with a one-minute interval between triggers to avoid immediate retriggering.

### Habitat types

Habitat types in the study area were determined using supervised habitat classification based on multispectral Landsat 5 TM imagery from 28 November 2010 (Klapproth 2010). The habitat classification distinguished six physiognomic habitat types (gallery forest/forest, savannah woodland, tree/shrub savannah, grass savannah, temporary wetlands, and wetlands) and focused on 158 km² in the Simenti region. We aggregated these six habitat types into three overall classes (gallery forest/forest = *forest*, savannah woodland, tree/shrub savannah, grass savannah = *savannah*, temporary wetlands, wetlands = *wetland*). Of our 37 camera-trap sampling points, 10 were in forest habitats, 25 in savannah habitats, and 2 in wetlands.

### Data analysis Global diversity

We uploaded all pictures to the platform “Agouti” for a first, AI-assisted, species identification (Casaer et al. 2019). We then reviewed all the observations annotated by the AI to confirm or modify the AI-assisted annotation. We sampled 13,161 camera-trap days between February 2022 and March 2023 (per sampling point: 217–390 days; mean = 355, SD = 37.3). We recorded a total of 52,585 events, containing 25,990 animal observations, 24,365 ‘blank’ observations (i.e., moving grass, bushfires), 732 vehicles (i.e., cars, planes, helicopters), 477 humans, and 1,021 unidentified observations (unidentifiable animals, or shapes that were not clearly animals or blank). We considered consecutive triggers as separate events if they involved a different species or occurred more than 120 seconds apart. In cases where a camera was clearly retriggered continuously by the same individual or group of individuals, we adjusted events manually.

### Species accumulation curves

To assess whether our sampling effort adequately captured the species present at the study site, we calculated species accumulation curves for both time and space using the *vegan* package version 2.7-2 (Oksanen et al. 2025) in R version 4.4.2 (R Core Team 2024). We used the function *specaccum* with 1,000 random permutations (without replacement) of the observed data to estimate the mean accumulation curves and their associated variation. We further calculated temporal and spatial sampling coverage Q (Fagen & Goldman 1977) as follows:

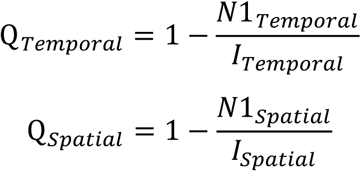

Here, N1_Temporal_ is the number of species recorded only once across all camera events, and I_Temporal_ is the total number of camera events. N1_Spatial_ is the number of species that only occurred at one sampling point of the grid, and I_Spatial_ is the sum of the number of sites each species was detected. Values of Q close to 1 indicate high sampling completeness and a low probability of detecting additional species with further temporal (Q_Temporal_) or spatial (Q_Spatial_) coverage.

### Local diversity

To assess whether there are areas or habitat types of particular importance within the study site, we calculated local detection frequencies for each species corrected for effort across all 37 sampling points. For each sampling point, we determined the local species count (S) and the total number of animal sighting events per day. We then calculated the evenness (J) for each sampling point as 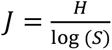, where *H* is the Shannon diversity index, and *S* is the total number of species detected at a given site. To further characterise the spatial variation in species occurrence, we calculated the naïve occupancy, i.e., the proportion of sampling sites at which a species was detected, for each species as an indicator of its spatial distribution across the study area. We then quantified the detection frequency for each species across the three habitat types within the camera-trapping grid to assess patterns of habitat use.

To assess differences in animal sighting rates and diversity measures between the three habitat types, we fitted separate models for each response variable (species count, animal sighting rate, evenness) with habitat type as the predictor using R. For species count and animal sighting rates, we fitted two generalized linear models with a negative binomial error structure and log link function (McCullagh & Nelder 1989) using the function *glmmTMB* of the *glmmTMB* package version 1.1.14 (Brooks et al. 2017). For animal-sighting rates, we included the log of the sampling effort per site as an offset term. For evenness, we fitted a generalized linear model with a beta error structure and logit link function (McCullagh & Nelder 1989) using the *glmmTMB* package version 1.1.14 (Brooks et al. 2017). For all models, we checked whether the assumption of no overdispersion was met. With dispersion parameters of 1.08, 1.105, and 1.117, respectively, the responses were not overdispersed given the models. To determine the significance of the predictors and to avoid ’cryptic multiple testing’ (Forstmeier & Schielzeth 2011), we used the function *drop1*, which drops each fixed effects predictor from the full model, one at a time, and conducts a likelihood ratio test comparing the resulting model with the full model (Dobson 2002).

For all models, we assessed stability by dropping each sampling point from the data one at a time and comparing the estimates from models fitted to these subsets to those from the full dataset. All models were of good stability (see results). We obtained confidence intervals of model estimates and fitted values using parametric bootstrapping (N=1000; function *simulate* of the package *glmmTMB*). To assess potential spatial autocorrelation in our data, we plotted the difference in residuals against the spatial distance between two sampling sites. This allowed us to visually evaluate whether residuals at spatially closer sites were related, indicating spatial autocorrelation. We further assessed Pearson correlation coefficients between residual difference and spatial distance. Neither the visual inspection nor the Pearson correlation coefficients indicated strong spatial autocorrelation in the data (Figure S1).

To further examine potential seasonal differences in habitat use, we classified all sightings from 1 June to 31 October as “wet season”, and from 1 November to 31 May as “dry season”. For each species, we then calculated the relative detection frequency per season and habitat type. Because the number of detections per season and habitat type varied substantially across species, we refrained from formally modelling their effects. Such models would have been underpowered and likely unreliable. Instead, we present potential patterns descriptively as exploratory results that may guide future research.

### Activity patterns

For species with 25 or more records across the camera-trapping grid, we quantified diel activity patterns from camera-trap detections using the *activity* package version 1.3.4 (Rowcliffe et al. 2014, Rowcliffe 2019) in R version 4.4.2 (R Core Team 2024). For each species, we extracted the timestamps of independent detections, converted them to radians, and fitted nonparametric circular kernel density functions to estimate activity distributions. We derived average sunrise and sunset times at the study site using the *suncalc package* (Thieurmel et al. 2019). We transformed clock times to solar times to account for seasonal shifts in day length and uneven sampling across dates, using the average-anchoring approach of Vázquez et al. (2019). Thus, activity is expressed relative to average sunrise and sunset times across the study period, weighted by the number of records per day. We then used the function *densityPlot* from the package *overlap* version 0.3.9 (Ridout & Linkie 2009) to visualise diel activity patterns for each species. To further investigate whether species use the three habitat types differently throughout the day, we estimated each species’ activity distribution for each habitat type separately.

## Results

### Global diversity

In total, we recorded 37 medium to large mammal species in the study area, including several of conservation concern according to the IUCN Red List (2026) (Table 1).

**Table 1:**
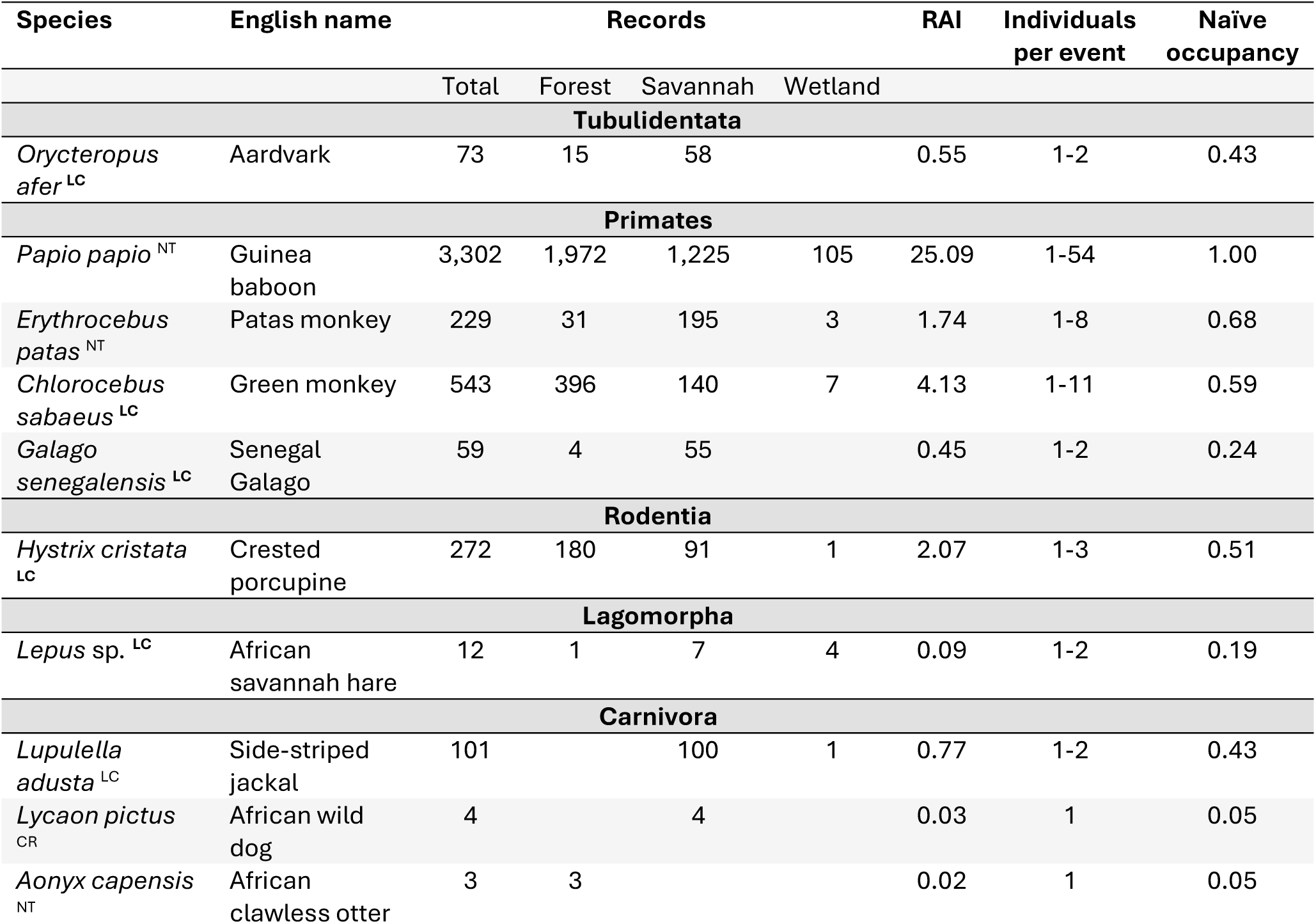
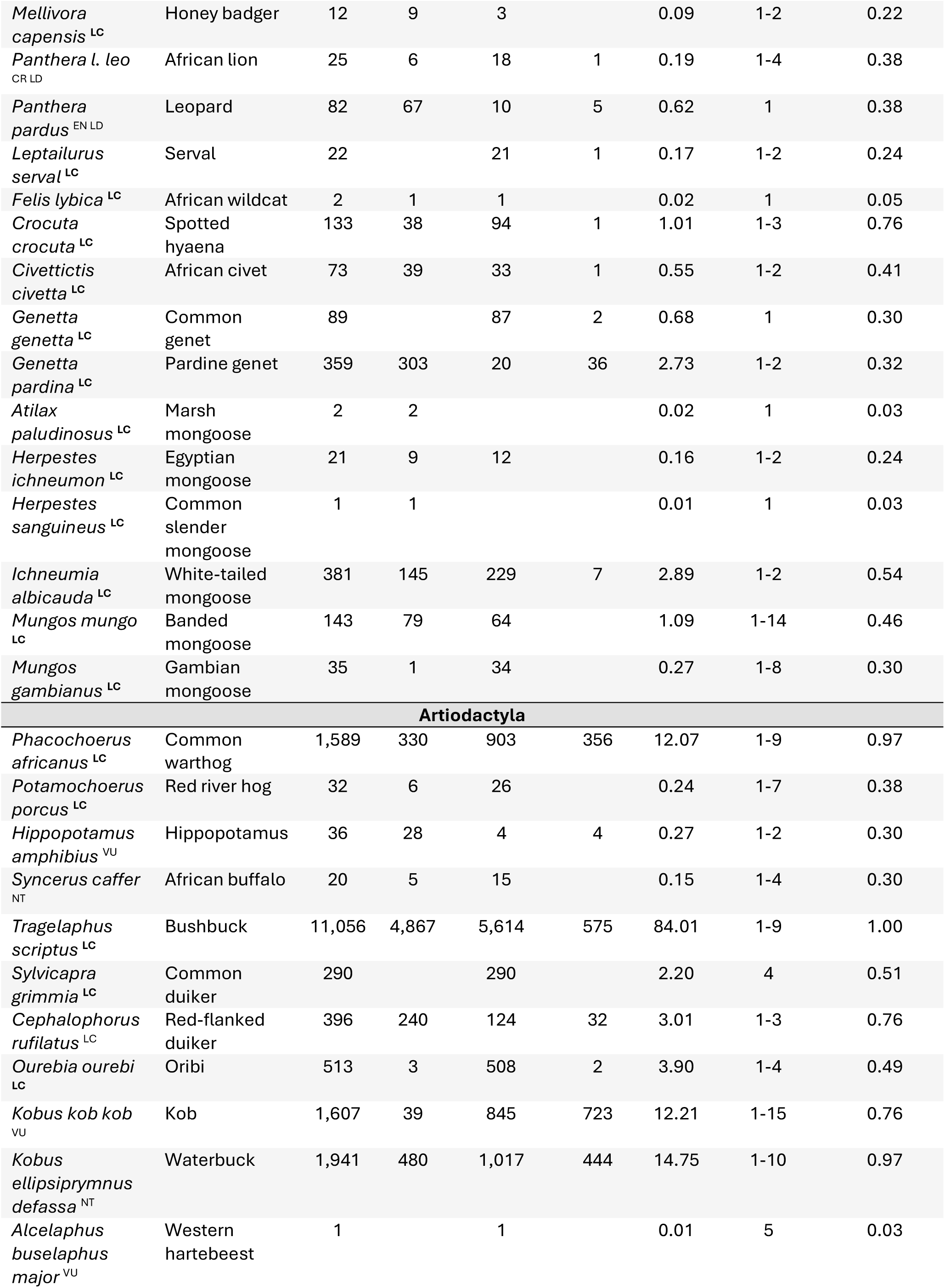
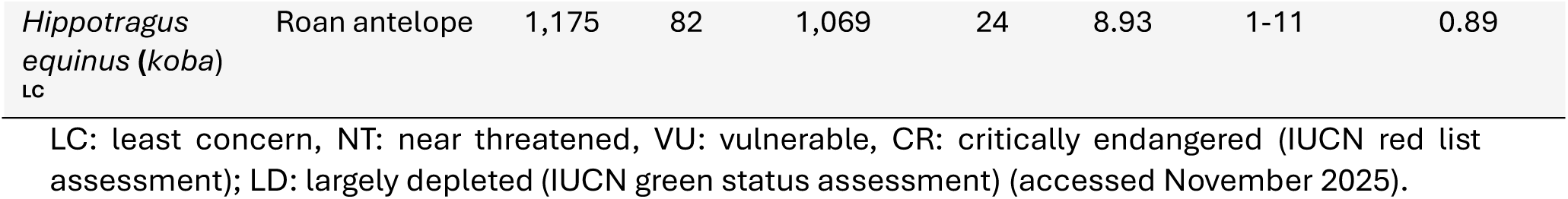
Overview of species identified from February 2022 to March 2023 in Simenti, Senegal, across 37 sampling points and 13,161 camera-trap days. Indicated is the total number of records, the number of records per habitat type, the relative abundance index (RAI) per 100 camera-trap days across the entire grid, the range of the number of individuals detected per animal sighting event, and the proportion of locations at which a species was recorded (naïve occupancy) across the camera-trapping grid. The IUCN Conservation status (IUCN 2026) for each species is included as a superscript.

| Species | English name | Records |  |  |  | RAI | Individuals<br>per event | Naïve<br>occupancy |
| --- | --- | --- | --- | --- | --- | --- | --- | --- |
|  |  | Total | Forest | Savannah | Wetland |  |  |  |
| Tubulidentata |  |  |  |  |  |  |  |  |
| *Orycteropus*<br>*afer*<sup>LC</sup> | Aardvark | 73 | 15 | 58 |  | 0.55 | 1-2 | 0.43 |
| Primates |  |  |  |  |  |  |  |  |
| *Papio papio*<sup>NT</sup> | Guinea<br>baboon | 3,302 | 1,972 | 1,225 | 105 | 25.09 | 1-54 | 1.00 |
| *Erythrocebus*<br>*patas*<sup>NT</sup> | Patas monkey | 229 | 31 | 195 | 3 | 1.74 | 1-8 | 0.68 |
| *Chlorocebus*<br>*sabaeus*<sup>LC</sup> | Green monkey | 543 | 396 | 140 | 7 | 4.13 | 1-11 | 0.59 |
| *Galago*<br>*senegalensis*<sup>LC</sup> | Senegal<br>Galago | 59 | 4 | 55 |  | 0.45 | 1-2 | 0.24 |
| Rodentia |  |  |  |  |  |  |  |  |
| *Hystrix cristata*<sup>LC</sup> | Crested<br>porcupine | 272 | 180 | 91 | 1 | 2.07 | 1-3 | 0.51 |
| Lagomorpha |  |  |  |  |  |  |  |  |
| *Lepus* sp. <sup>LC</sup> | African<br>savannah hare | 12 | 1 | 7 | 4 | 0.09 | 1-2 | 0.19 |
| Carnivora |  |  |  |  |  |  |  |  |
| *Lupulella*<br>*adusta*<sup>LC</sup> | Side-striped<br>jackal | 101 |  | 100 | 1 | 0.77 | 1-2 | 0.43 |
| *Lycaon pictus*<sup>CR</sup> | African wild<br>dog | 4 |  | 4 |  | 0.03 | 1 | 0.05 |
| *Aonyx capensis*<sup>NT</sup> | African<br>clawless otter | 3 | 3 |  |  | 0.02 | 1 | 0.05 |
| *Mellivora capensis* <sup>LC</sup> | Honey badger | 12 | 9 | 3 |  | 0.09 | 1-2 | 0.22 |
| *Panthera l. leo* <sup>CR LD</sup> | African lion | 25 | 6 | 18 | 1 | 0.19 | 1-4 | 0.38 |
| *Panthera pardus* <sup>EN LD</sup> | Leopard | 82 | 67 | 10 | 5 | 0.62 | 1 | 0.38 |
| *Leptailurus serval* <sup>LC</sup> | Serval | 22 |  | 21 | 1 | 0.17 | 1-2 | 0.24 |
| *Felis lybica* <sup>LC</sup> | African wildcat | 2 | 1 | 1 |  | 0.02 | 1 | 0.05 |
| *Crocuta crocuta* <sup>LC</sup> | Spotted hyaena | 133 | 38 | 94 | 1 | 1.01 | 1-3 | 0.76 |
| *Civettictis civetta* <sup>LC</sup> | African civet | 73 | 39 | 33 | 1 | 0.55 | 1-2 | 0.41 |
| *Genetta genetta* <sup>LC</sup> | Common genet | 89 |  | 87 | 2 | 0.68 | 1 | 0.30 |
| *Genetta pardina* <sup>LC</sup> | Pardine genet | 359 | 303 | 20 | 36 | 2.73 | 1-2 | 0.32 |
| *Atilax paludinosus* <sup>LC</sup> | Marsh mongoose | 2 | 2 |  |  | 0.02 | 1 | 0.03 |
| *Herpestes ichneumon* <sup>LC</sup> | Egyptian mongoose | 21 | 9 | 12 |  | 0.16 | 1-2 | 0.24 |
| *Herpestes sanguineus* <sup>LC</sup> | Common slender mongoose | 1 | 1 |  |  | 0.01 | 1 | 0.03 |
| *Ichneumia albicauda* <sup>LC</sup> | White-tailed mongoose | 381 | 145 | 229 | 7 | 2.89 | 1-2 | 0.54 |
| *Mungos mungo* <sup>LC</sup> | Banded mongoose | 143 | 79 | 64 |  | 1.09 | 1-14 | 0.46 |
| *Mungos gambianus* <sup>LC</sup> | Gambian mongoose | 35 | 1 | 34 |  | 0.27 | 1-8 | 0.30 |
| **Artiodactyla** |  |  |  |  |  |  |  |  |
| *Phacochoerus africanus* <sup>LC</sup> | Common warthog | 1,589 | 330 | 903 | 356 | 12.07 | 1-9 | 0.97 |
| *Potamochoerus porcus* <sup>LC</sup> | Red river hog | 32 | 6 | 26 |  | 0.24 | 1-7 | 0.38 |
| *Hippopotamus amphibius* <sup>VU</sup> | Hippopotamus | 36 | 28 | 4 | 4 | 0.27 | 1-2 | 0.30 |
| *Syncerus caffer* <sup>NT</sup> | African buffalo | 20 | 5 | 15 |  | 0.15 | 1-4 | 0.30 |
| *Tragelaphus scriptus* <sup>LC</sup> | Bushbuck | 11,056 | 4,867 | 5,614 | 575 | 84.01 | 1-9 | 1.00 |
| *Sylvicapra grimmia* <sup>LC</sup> | Common duiker | 290 |  | 290 |  | 2.20 | 4 | 0.51 |
| *Cephalophorus rufilatus* <sup>LC</sup> | Red-flanked duiker | 396 | 240 | 124 | 32 | 3.01 | 1-3 | 0.76 |
| *Ourebia ourebi* <sup>LC</sup> | Oribi | 513 | 3 | 508 | 2 | 3.90 | 1-4 | 0.49 |
| *Kobus kob kob* <sup>VU</sup> | Kob | 1,607 | 39 | 845 | 723 | 12.21 | 1-15 | 0.76 |
| *Kobus ellipsiprymnus defassa* <sup>NT</sup> | Waterbuck | 1,941 | 480 | 1,017 | 444 | 14.75 | 1-10 | 0.97 |
| *Alcelaphus buselaphus major* <sup>VU</sup> | Western hartebeest | 1 |  | 1 |  | 0.01 | 5 | 0.03 |
| *Hippotragus equinus (koba)*<br>LC | Roan antelope | 1,175 | 82 | 1,069 | 24 | 8.93 | 1-11 | 0.89 |

LC: least concern, NT: near threatened, VU: vulnerable, CR: critically endangered (IUCN red list assessment); LD: largely depleted (IUCN green status assessment) (accessed November 2025).

### Species accumulation curves

Species accumulation curves based on camera-trap days and sampling points approached an asymptote (Figure 2). Values for Q were 1 or close to one for both temporal and spatial sampling coverage, indicating that the sampling effort was sufficient to capture most medium- to large-bodied species detectable by our setup.

**Figure 2:**
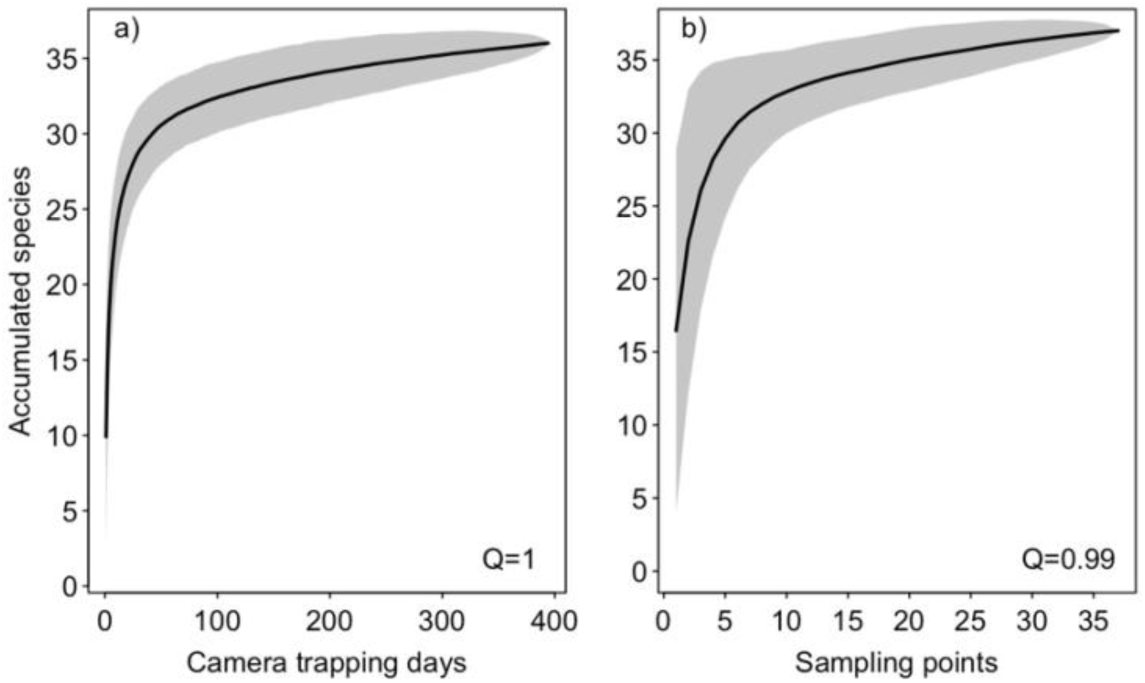
a) Temporal and b) spatial species accumulation curves. The black line depicts the estimated mean, and the grey shaded areas depict confidence intervals.

**Figure 3:**
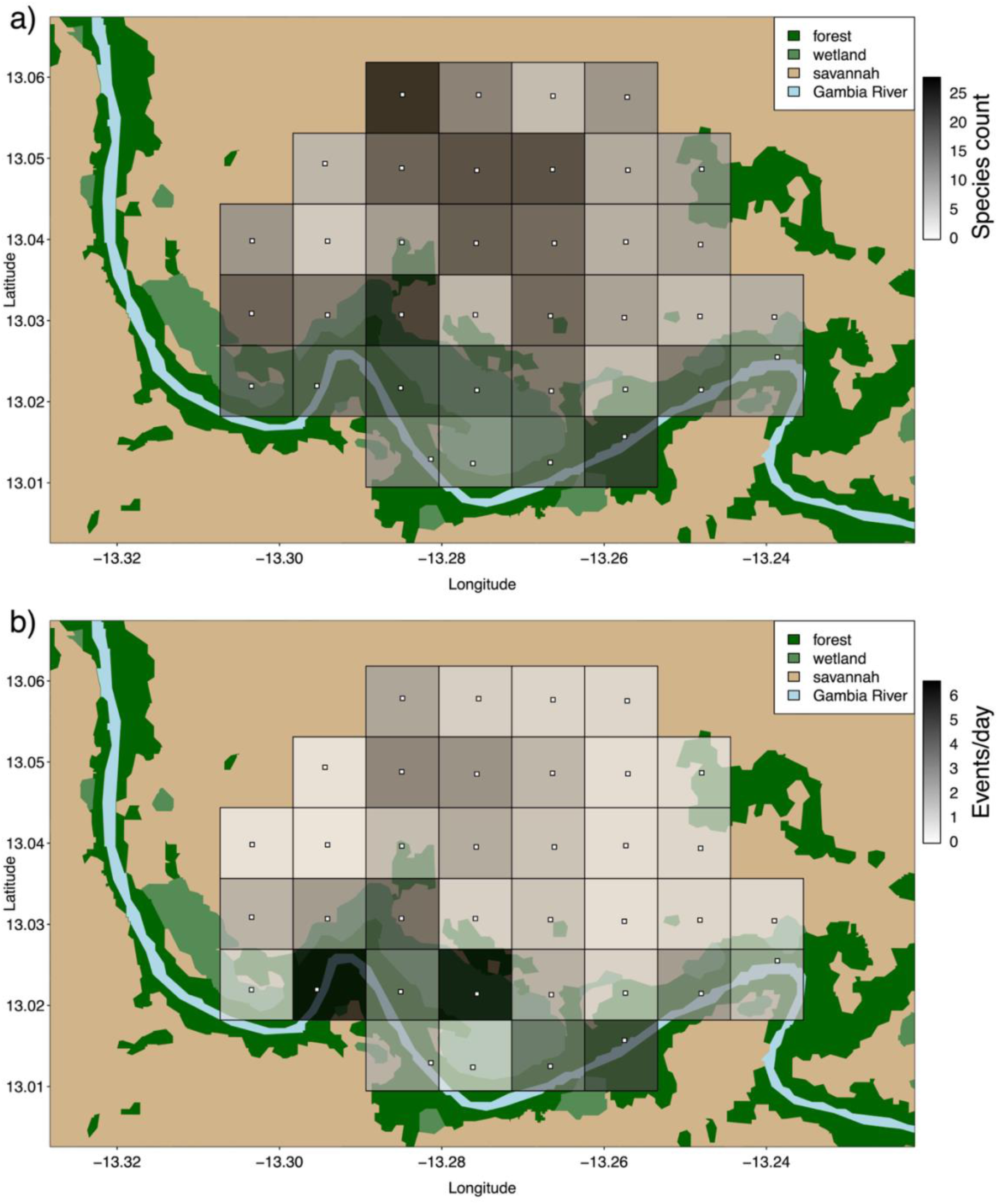
a) Local species counts and b) animal sighting events per day and sampling point (white squares), both from February 2022 to March 2023 across the study site.

### Local diversity

The number of species recorded per sampling point ranged from 5 to 28, with a median of 16 (Figure 3a). The number of animal-sighting events per day and per camera ranged from 0 to 6 (Figure 3b). Evenness values were moderate across the study site, ranging from 0.37 to 0.79 with a mean of 0.6, suggesting relatively uniform species distributions at most sites. The total number of species recorded did not differ between habitat types. Habitat type was a significant predictor of evenness across the study site (Table 2). Evenness values were significantly higher at savannah sites than at forest sites. Habitat type was also a significant predictor of the number of animal sightings per day. However, in contrast to evenness values, the number of animal sighting events per day was higher in forest habitats than in savannahs. Given the small number of wetland sampling sites (n=2), we do not interpret differences between wetlands and other habitat types.

**Table 2:**
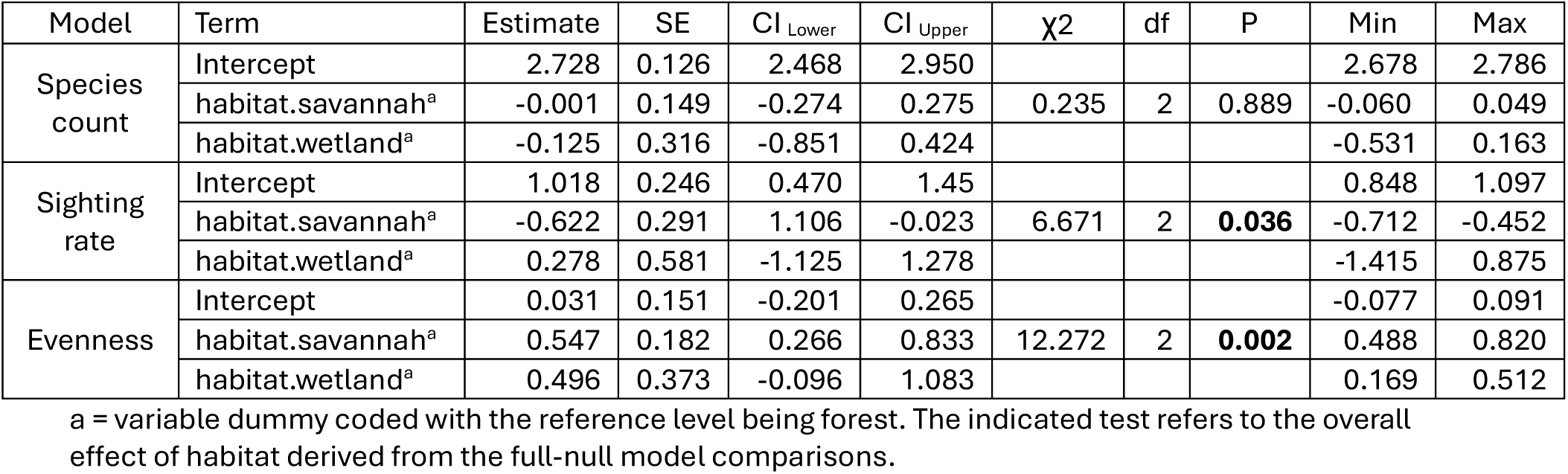
Model results on species count, animal sighting rates, and evenness; estimates, standard errors, confidence intervals, likelihood ratio tests, significance tests, and range of estimates derived from dropping each sampling site one at a time.

| Model | Term | Estimate | SE | CI Lower | CI Upper | $\chi^2$ | df | P | Min | Max |
| --- | --- | --- | --- | --- | --- | --- | --- | --- | --- | --- |
| Species count | Intercept | 2.728 | 0.126 | 2.468 | 2.950 |  |  |  | 2.678 | 2.786 |
|  | habitat.savannah<sup>a</sup> | -0.001 | 0.149 | -0.274 | 0.275 | 0.235 | 2 | 0.889 | -0.060 | 0.049 |
|  | habitat.wetland<sup>a</sup> | -0.125 | 0.316 | -0.851 | 0.424 |  |  |  | -0.531 | 0.163 |
| Sighting rate | Intercept | 1.018 | 0.246 | 0.470 | 1.45 |  |  |  | 0.848 | 1.097 |
|  | habitat.savannah<sup>a</sup> | -0.622 | 0.291 | 1.106 | -0.023 | 6.671 | 2 | **0.036** | -0.712 | -0.452 |
|  | habitat.wetland<sup>a</sup> | 0.278 | 0.581 | -1.125 | 1.278 |  |  |  | -1.415 | 0.875 |
| Evenness | Intercept | 0.031 | 0.151 | -0.201 | 0.265 |  |  |  | -0.077 | 0.091 |
|  | habitat.savannah<sup>a</sup> | 0.547 | 0.182 | 0.266 | 0.833 | 12.272 | 2 | **0.002** | 0.488 | 0.820 |
|  | habitat.wetland<sup>a</sup> | 0.496 | 0.373 | -0.096 | 1.083 |  |  |  | 0.169 | 0.512 |

<sup>a</sup> = variable dummy coded with the reference level being forest. The indicated test refers to the overall effect of habitat derived from the full-null model comparisons.

Naïve occupancy of species ranged from 0.03 to 1, where 0.03 indicates occurrence at only one out of 37 sampling sites, and 1 indicates occurrence at all sites. *Papio papio* and *Tragelaphus scriptus* were most widely distributed (1), followed by *Kobus ellipsiprymnus* and *Phacochoerus africanus* (0.97), and by *Hippotragus equinus* (0.89). Maps showing the number of individuals detected per species and sampling site within 100 days of camera-trapping are provided in the supplementary material (Figures S2-S38). Patterns of habitat use varied between species. Fifteen out of 37 species were mostly detected in savannah habitats (e.g., *Lupulella adusta*, *Genetta genetta*, *Sylvicapra grimmia*), while seven species were mostly detected in forests (e.g., *Papio papio*, *Panthera pardus*, *Genetta pardina*) (Table 1). The remaining species were either detected too infrequently or showed no clear pattern of habitat use.

We observed varying patterns of seasonal habitat use across species. We only report seasonal differences in habitat use for species with at least 25 total records. For some species (e.g., *Papio papio*, *Chlorocebus sabaeus*, *Hystrix cristata*, *Mungos mungo*), habitat use did not differ between seasons. Eight out of 26 species showed a shift from forests in the dry season to savannahs in the wet season (e.g., *Cephalophorus rufilatus*, *Civettictis civetta*, *Erythrocebus patas*, *Panthera pardus*). *Kobus ellipsiprymnus* seemed to shift from forests in the dry season to wetlands in the wet season. *K. kob* shifted from wetlands in the dry season to savannahs in the wet season. Lastly, *Phacochoerus africanus* shifted from savannahs in the dry season to wetlands in the wet season (Table S1).

### Activity patterns

For 26 out of 37 detected mammal species, we were able to estimate diel activity patterns. Some species exhibited clearly nocturnal (e.g., *Hystrix cristata, Orycteropus afer*) (Figure 5), crepuscular (e.g., *Lupulella adusta, Panthera pardus*) (Figure 7a, b), or diurnal (e.g., *Papio papio, Phacochoerus africanus*) (Figures 6 & 8a) activity patterns. For other species, particularly ungulates (e.g., *Tragelaphus scriptus*, *Hippotragus equinus*) (Figure 8b), diel activity patterns were more ambiguous, showing multiple peaks across day and night and a relatively high baseline of nocturnal activity.

**Figure 4:**
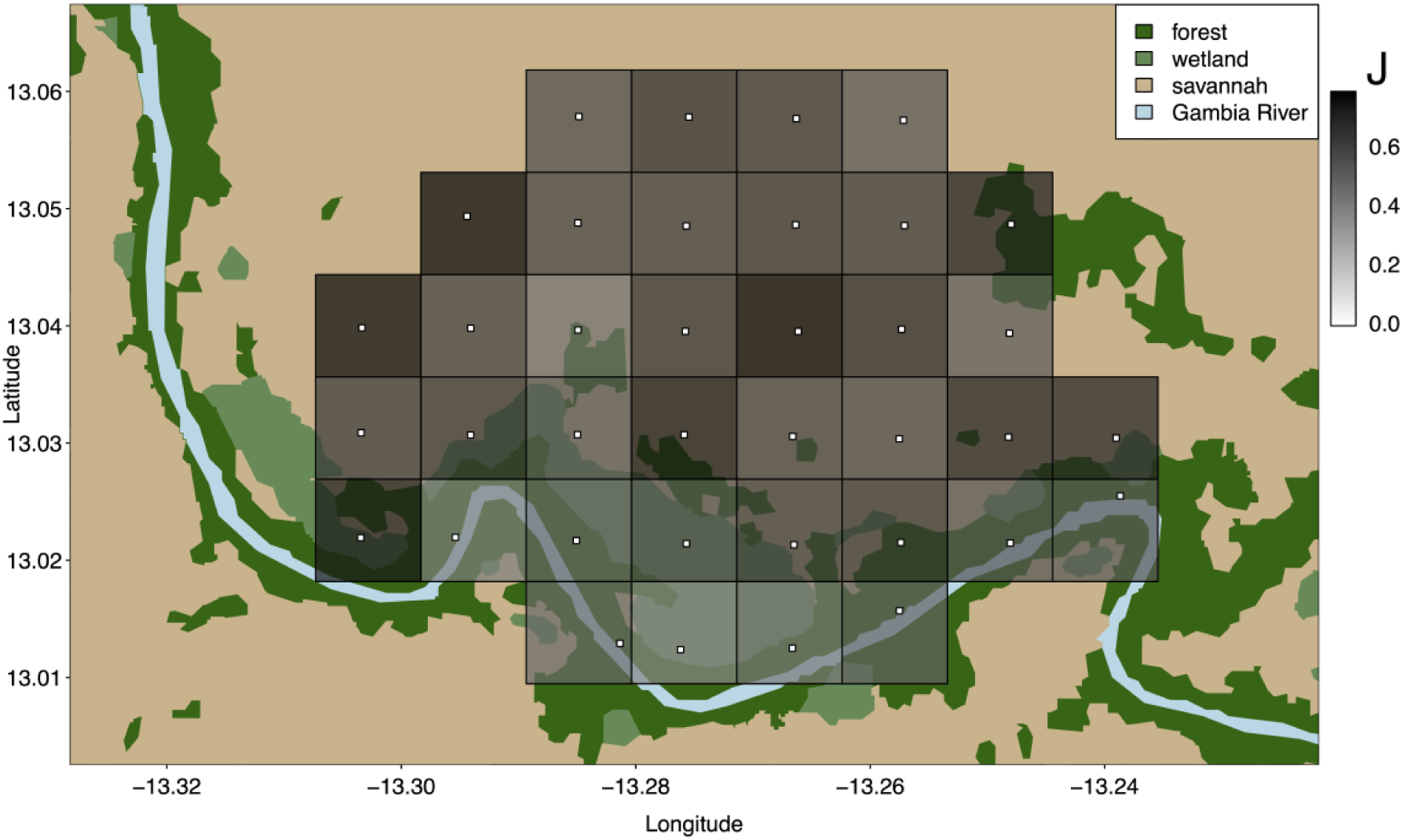
Local evenness values (J) calculated per sampling point (white squares) from February 2022 to March 2023 across the study site.

**Figure 5:**
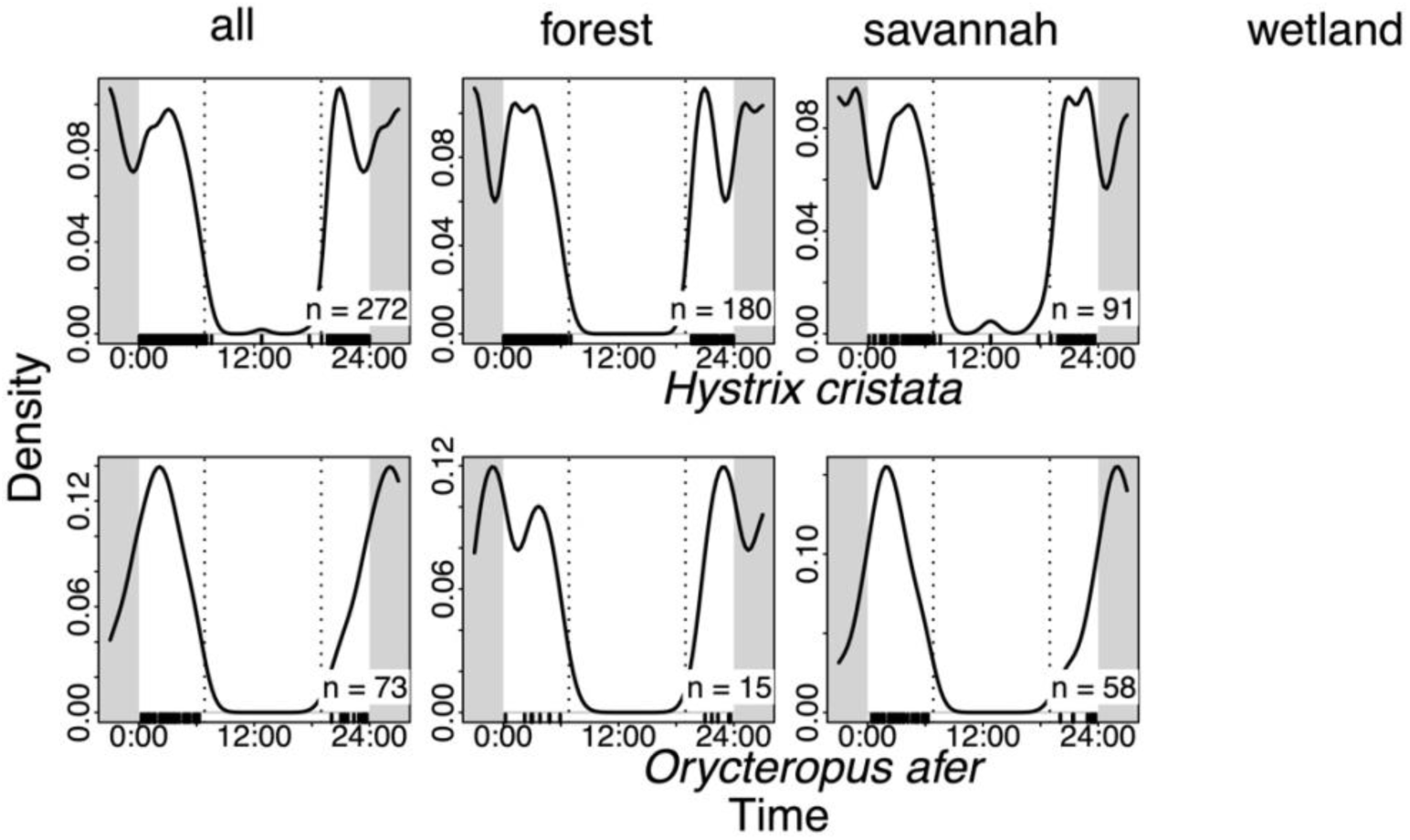
Diel activity patterns for *Hystrix cristata* and *Orycteropus afer* and respective sample sizes (n) across the study site (all) and by habitat type (forest, savannah, wetland). Note that blank panels in the plot are due to insufficient data coverage (only one or no sighting of a species in a given habitat type). Dashed lines indicate the average times of sunrise and sunset at the study site. The original observations are displayed as a rug along the timeline at the bottom of the plots.

**Figure 6:**
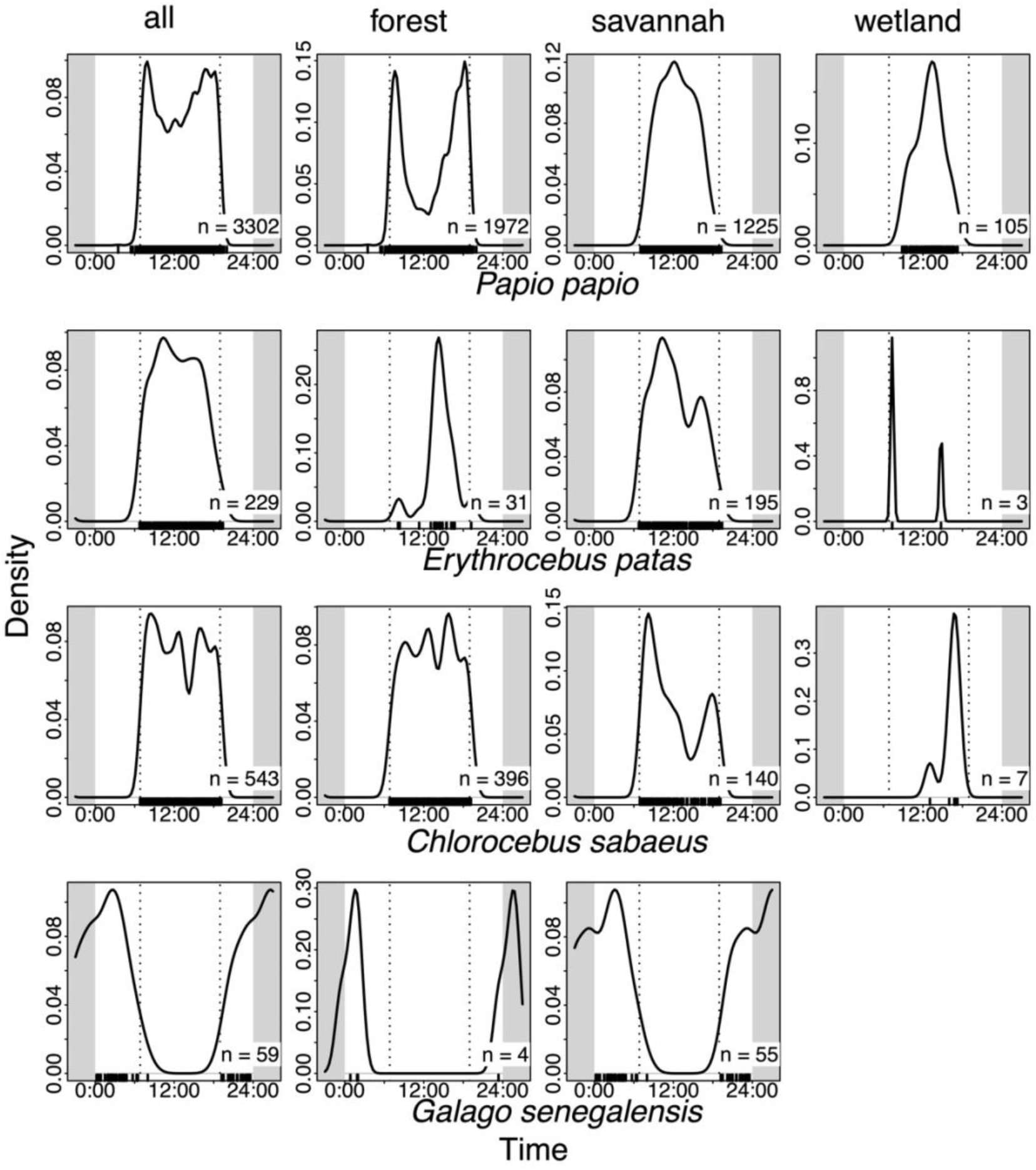
Diel activity patterns for the recorded primate species and respective sample sizes (n) across the study site (all) and by habitat type (forest, savannah, wetland). Note that blank panels in the plot are due to insufficient data coverage (only one or no sighting of a species in a given habitat type). Dashed lines indicate the average times of sunrise and sunset at the study site. The original observations are displayed as a rug along the timeline at the bottom of the plots.

**Figure 7a:**
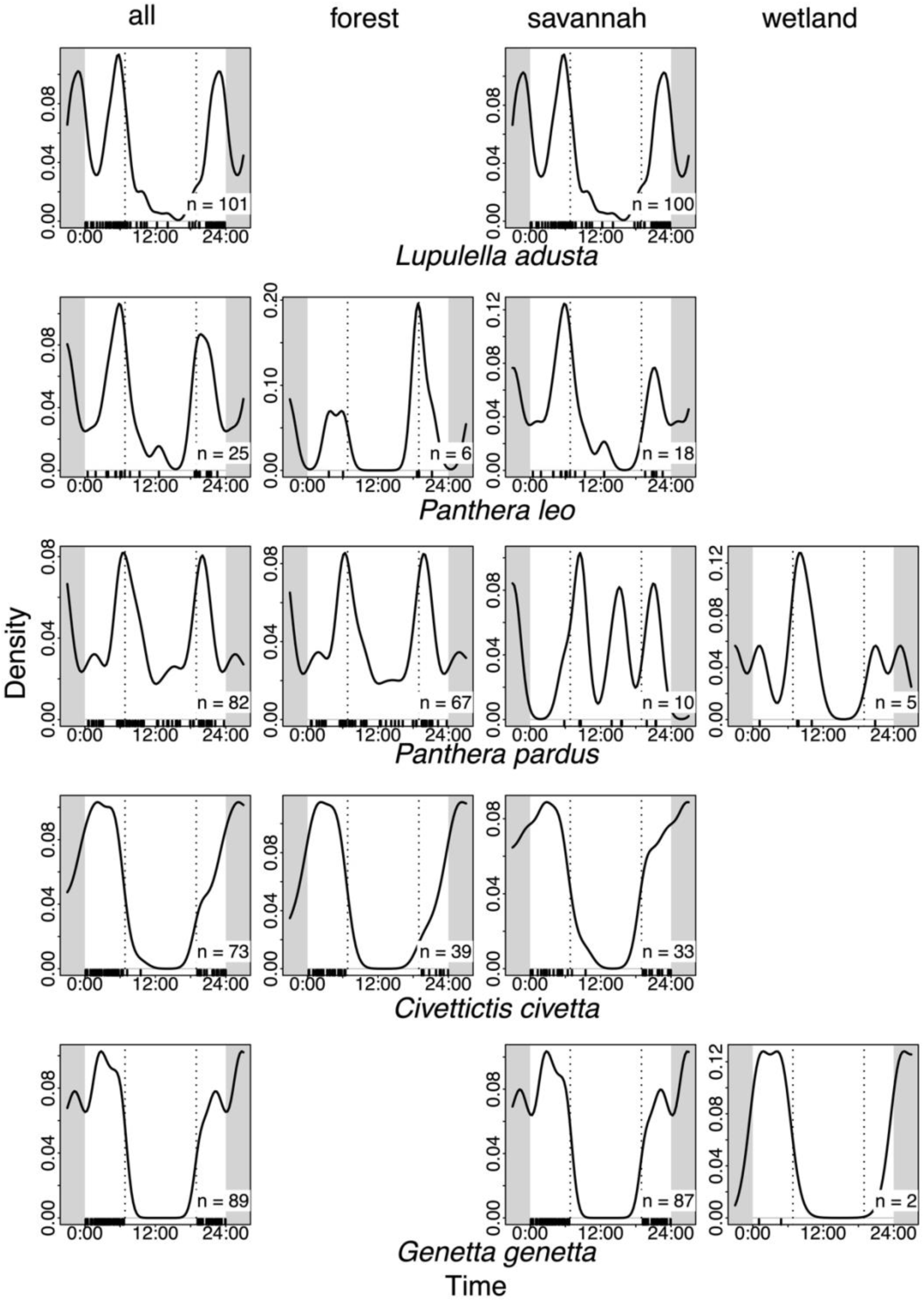
Diel activity patterns of recorded carnivore species and respective sample

**Figure 7b:**
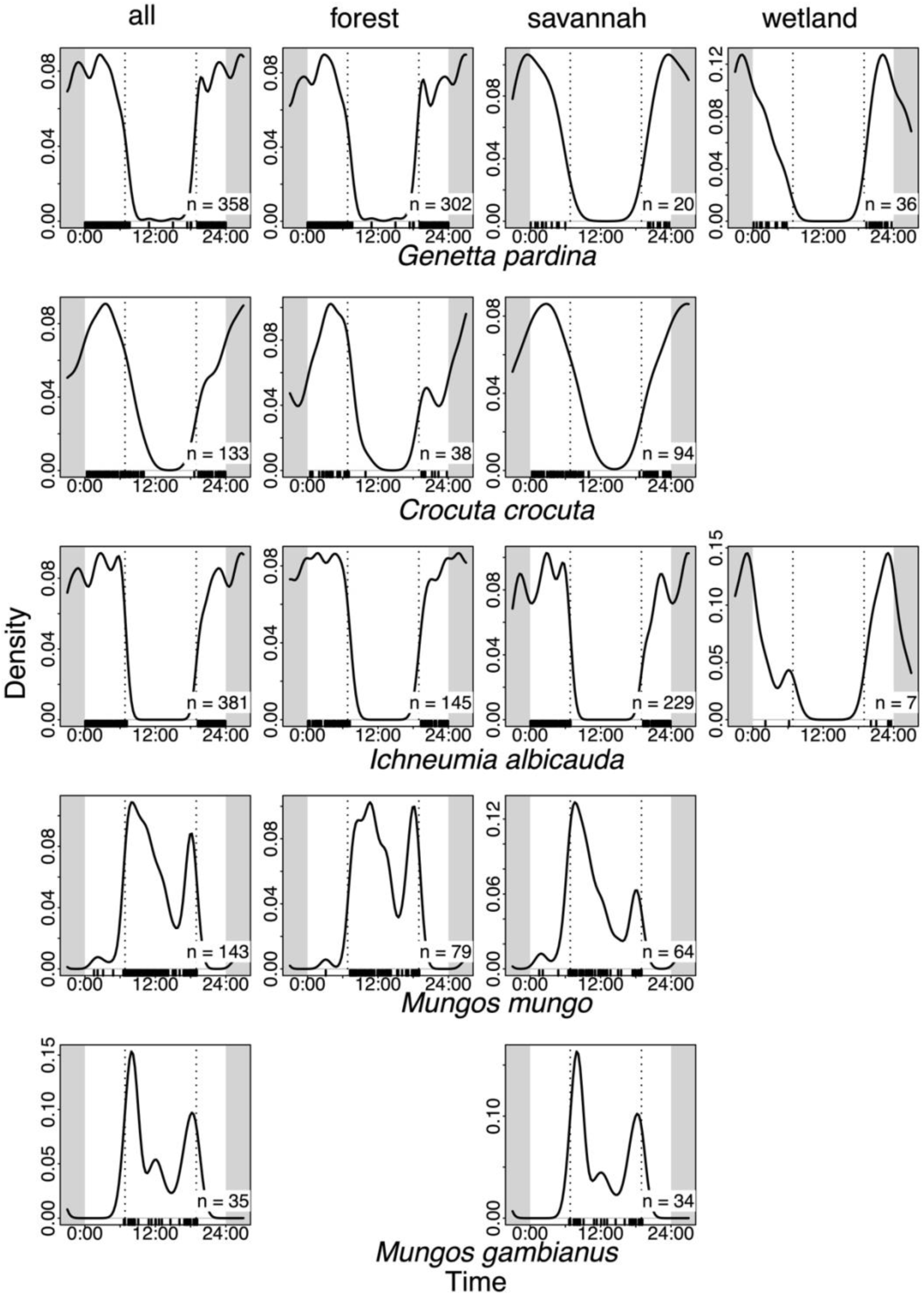
Diel activity patterns of recorded carnivore species and respective sample

**Figure 8a:**
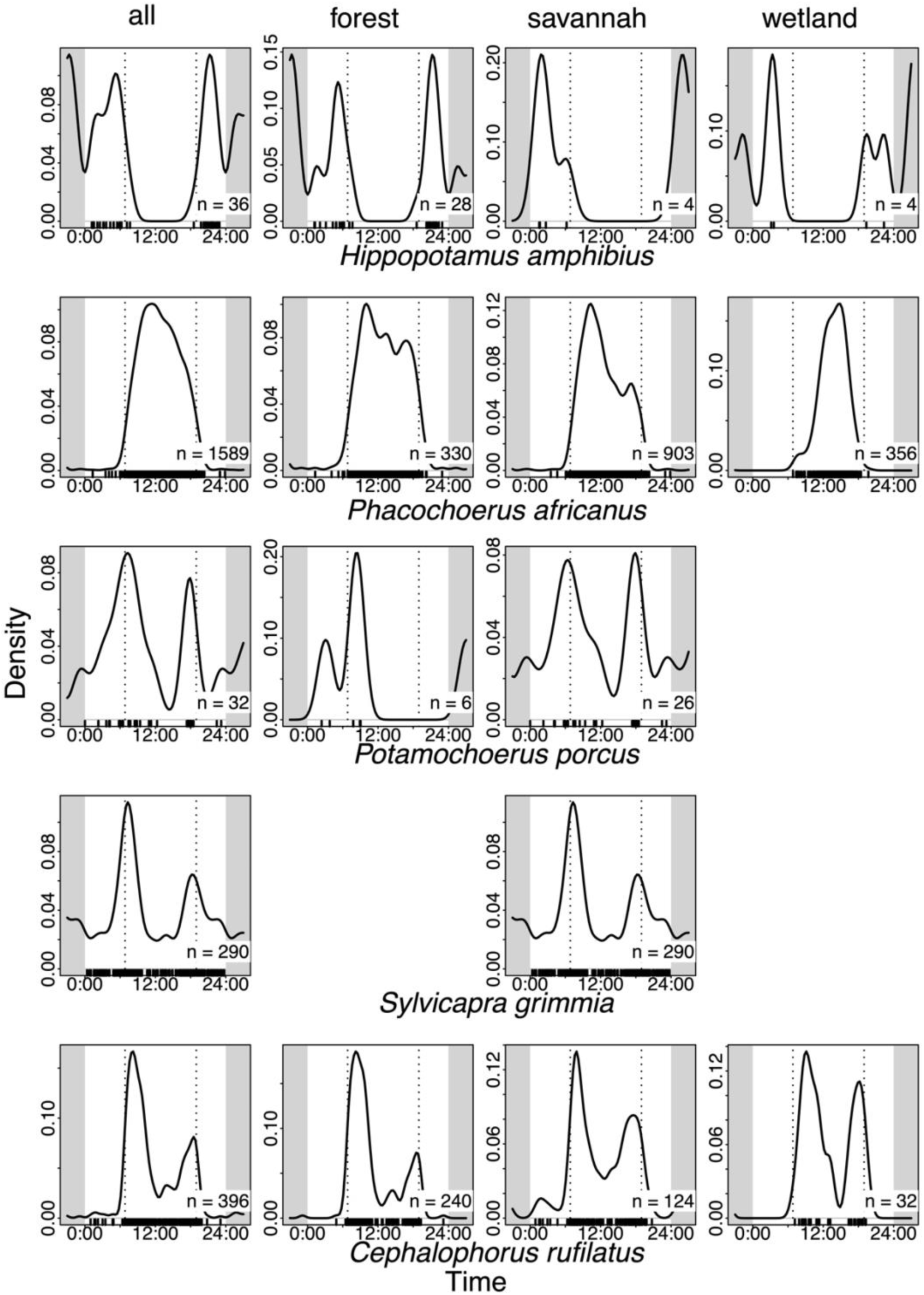
Diel activity patterns of recorded artiodactyl species and respective sample

**Figure 8b:**
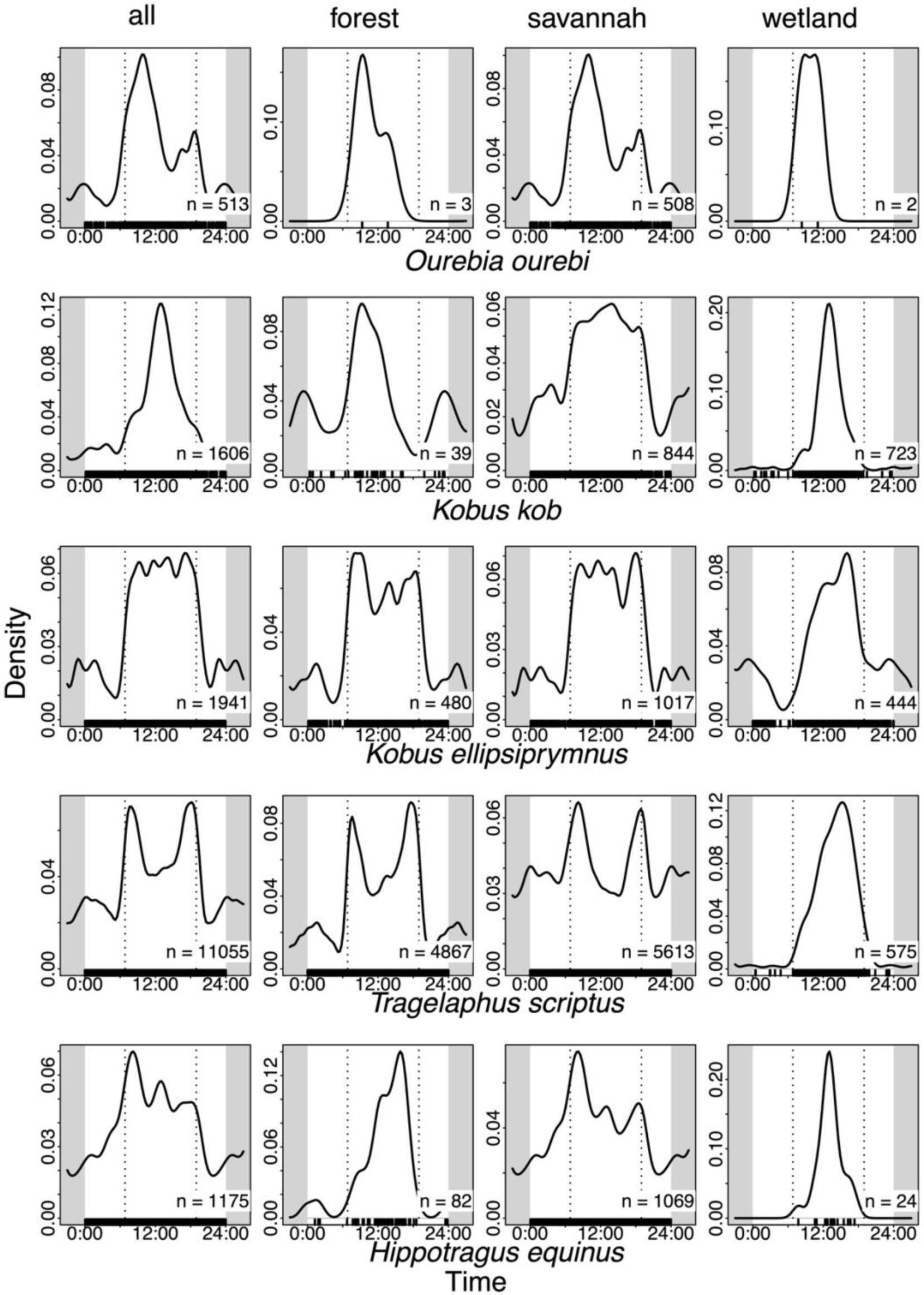
Diel activity patterns of recorded artiodactyl species and respective sample sizes (n) across the study site (all) and by habitat type (forest, savannah, wetland). Dashed lines indicate the average times of sunrise and sunset at the study site. The original observations are displayed as a rug along the timeline at the bottom of the plots.

Most species’ activity distribution did not differ between habitat types (Figures 5-8b). *Papio papio* exhibited higher levels of activity in forest habitats in the mornings and evenings, and in savannahs and wetlands during the day. Some ungulates (*Kobus kob*, *Tragelaphus scriptus*, *Hippotragus equinus*) seemed to shift their activity from wetlands during the day to forests and savannahs during the night. Although very low sample sizes did not permit a reliable estimation of activity distribution, we included these panels to visualize data availability and distribution, but refrained from interpreting any patterns.

## Discussion

### Global diversity

Our camera-trap survey provides a comprehensive inventory of medium- to large-sized terrestrial mammals in Niokolo-Koba National Park. We documented 37 species, representing the majority of medium-sized to large mammal species expected to occur in the Simenti area based on IUCN range maps (IUCN 2026). When considering only medium- to large-sized mammals, our inventory largely aligns with both historical and more contemporary records for the region (Dupuy 1971, Kane 2014, Rabeil et al. 2018, Horion et al. 2024, Mirghani et al. 2025). Although small, arboreal, or aerial species fall outside the detection range of our setup, our study effectively captured the local terrestrial mammal assemblage, including several species of conservation concern.

Several of the park’s flagship species, including *Pan troglodytes verus* and *Taurotragus derbianus*, were not recorded in our study area, although they occur in other sectors of the park (Kane 2014, Rabeil et al. 2018, Horion et al. 2024). Their absence may reflect a combination of factors, including limited suitable habitat, floral and structural diversity, and the presence of large predators at our study site (Lindshield et al. 2019, Wessling et al. 2020, Stehlíková 2023). Other taxa expected to occur at low densities throughout the park, including caracals (*Caracal caracal*), Bohor reedbuck (*Redunca redunca*), African palm civet (*Nandinia binotata*), and rock hyraxes (*Procavia ruficeps*), were also not recorded, but have been reported to occur in the southeastern sections of the park and surrounding areas (Kane 2014, Horion et al. 2024, Mirghani et al. 2025). Their absence may likewise reflect local ecological conditions, such as the absence of rocky cliffs or other habitat features important for these species.

Extremely rare species, such as *Loxodonta* sp. and giant pangolin (*Smutsia gigantea*), were absent from our dataset. Recent records suggest that *Loxodonta* sp. may be limited to a single individual in the southern sector of the park (Panthera 2025), while the detection of *S. gigantea* in 2023 represents the first confirmed observation in 24 years (Ndiaye et al. 2024). These two species do not seem to occur in the Simenti area.

The Relative Abundance Index (RAI) values for most species at our study site are broadly comparable to those reported by Kane (2014), though with some notable differences. Carnivores (e.g., *Panthera leo, P. pardus*) were detected up to 10 times more frequently in Kane’s study. This is likely due to differences in sampling design. While Kane (2014) deliberately selected an area of particularly high large-carnivore abundance identified in a pilot study, our camera-trapping grid was designed to systematically cover the entire study area for the duration of a year.

### Local diversity

Our analyses revealed that evenness was higher in savannah than in forest habitats. However, the total number of species detected did not differ between habitat types. The number of animal sighting events per day was higher in forests than in savannah habitats. While detection rates at wetland sites were relatively high, direct comparisons with other habitats would be misleading, as wetlands were underrepresented in our setup and strongly affected by seasonal flooding and greater visibility. Nevertheless, wetlands and the Gambia River are the only perennial water sources in the area and are likely important landscape features for many species, particularly during the dry season (UNESCO World Heritage Centre 2025). Elevated animal sighting rates at some forest sites near the Gambia River, in contrast, appear to be driven by a subset of species disproportionately using these sites. In particular, certain parts of the gallery forest along the Gambia River exhibited this pattern, likely because Guinea baboons used tall trees as sleeping sites (Zinner et al. 2021, Ohrndorf et al. 2025b). Taken together, these patterns suggest that the landscape as a whole supports consistently high local diversity, but not all species use it homogeneously.

Several of the most frequently detected species (e.g., *Tragelaphus scriptus, Papio papio, Phacochoerus africanus, Kobus ellipsiprymnus, Hippotragus equinus*) were recorded across the entire study site, suggesting extensive use of the available habitat, whether due to high abundance, wide spatial movements, large home ranges, or a combination thereof. Other species showed more pronounced spatial clustering, likely reflecting habitat preferences or localised activity centres, potentially constrained by smaller home ranges or territories (e.g., *Galago senegalensis, Genetta pardina, Ourebia ourebi*).

In our descriptive assessment of habitat-specific sighting frequencies, many observed species shifted from forests in the dry season to savannahs in the wet season. This pattern may reflect a strategy to reduce exposure to sun and heat during the hottest months of the year, and to remain close to surface water for drinking (Brain & Mitchell 1999, Terrien et al. 2011, Boyers et al. 2019). Gallery forests along the Gambia River and the Mare de Simenti provide access to the only perennial water sources. For grazers and folivores, and potentially for frugivores, omnivores, and insectivores, the shift toward savannah habitats in the wet season may also be driven by food availability. At the study site, leaves and fruits of many trees, as well as invertebrates and much of the palatable herbaceous vegetation in savannah habitats, typically only become available after the first rains of the wet season (Ohrndorf et al. 2025a).

*Kobus ellipsiprymnus* and *Phacochoerus africanus* showed a shift towards wetlands in the wet season. Both species are well adapted to marshy soils and often seek out partially flooded areas with waterlogged substrates (Cumming 1975, Kingdon et al. 2013). Wetlands may also provide better access to drinking water, wallows, and higher food availability, with softer soils that allow digging for roots and below-ground storage organs (Cumming 1975, Kingdon et al. 2013). In contrast, *Kobus kob* shifted from wetlands to savannah habitats during the wet season. Although still highly dependent on surface water, this species typically prefers drier grasslands (Kingdon et al. 2013, Antwi et al. 2018). During the wet season, the increased availability of surface water and palatable grasses in savannahs likely allows them to use these habitats more frequently.

### Activity patterns

The temporal activity patterns estimated from our dataset largely reflect the expected diel behavioural patterns of the recorded species. Many species were clearly assignable to predominantly diurnal, nocturnal, or crepuscular activity patterns, indicating well-defined temporal niches. Some diurnal species (e.g., *Mungos mungo*, *M. gambianus*, *Papio papio*) showed a noticeable decline in detections during midday and early afternoon hours, potentially reflecting behavioural thermoregulation in response to high temperatures, particularly in open habitats (Dzingwena et al. 2025).

For other species, the categorisation of temporal activity was less distinct. Many ungulate species (e.g., *Tragelaphus scriptus*, *Hippotragus equinus*, *Kobus ellipsiprymnus*) exhibited moderate to high levels of activity throughout the 24-hour cycle, consistent with broad species-level descriptions reporting flexible activity patterns across the day (Kingdon et al. 2013). Some species deviated slightly from expectations. *Panthera leo* exhibited a more crepuscular activity pattern than reported for the region (Gueye et al. 2024, Horion et al. 2024), though this might be influenced by the small number of lion detections in our dataset. *Panthera pardus* and *Lupulella adusta* showed more activity during the day than commonly reported, a pattern often associated with low levels of anthropogenic disturbance (Kingdon et al. 2013, Gaynor et al. 2018).

When examining species’ activity distributions across the three habitat types separately, we found that some species shifted their habitat use throughout the day. *Papio papio*, in particular, showed a pronounced bimodal pattern of forest use in the early mornings and late afternoons to evenings, with increased use of savannah and wetland habitats during the day. This pattern likely reflects the availability of suitable sleeping trees in gallery forests, which are heavily used by baboons, as well as the spatial segregation of sleeping sites and most daytime foraging areas (Zinner et al. 2021, Ohrndorf et al. 2025a, b). *Kobus kob*, *Tragelaphus scriptus*, and *Hippotragus equinus* used wetland habitats mostly during the day, shifting to savannah and forest habitats at night. This shift may be driven by a preference for habitats offering greater cover during periods of increased vulnerability to predation (Bonnot et al. 2013, Burkepile et al. 2013, Anderson et al. 2016). Most recorded species, however, did not show a clear pattern of habitat use across the day, or lacked sufficient data to reliably assess habitat-specific activity distributions.

## Conclusion

Our camera-trap survey provides high-resolution, up-to-date information on the community of medium-sized to large mammals in the Simenti region of NKNP, documenting a species assemblage broadly consistent with historical and contemporary records. Evenness values were moderate to high across the landscape, and even areas with lower diversity appeared to be heavily used by certain species, as indicated by elevated animal sighting rates. The recorded species appeared to use the local habitat mosaic relatively uniformly, highlighting the ecological importance of all habitat types. A descriptive assessment of habitat-specific sighting frequencies suggests seasonal shifts in habitat use for several species, likely indicating strategies for thermoregulation and/or varying levels of food availability across habitats and seasons. Diel activity patterns were consistent with expectations for the observed species. Some species exhibited differential use of the available habitats throughout the day. Altogether, our results provide a useful reference for future studies and monitoring efforts and help characterise the local species assemblage of the Simenti area within the larger NKNP ecosystem.

## Declarations

### Authors’ contributions

LO, JF and DZ designed the study. LO, ND, IGD, DC, CYKC and ABD set up and maintained the camera-trapping grid from February 2022 to March 2023. LO and AB annotated the camera-trap imagery. LO, AMZ and AB analysed the data and prepared the figures. LO prepared the manuscript with contributions from all co-authors. All authors read, edited and approved the final manuscript.

## Acknowledgements

We are grateful to the Direction des Parcs Nationaux (DPN) and the Ministère de l’Environnement et de la Protéction de la Nature (MEPN) de la République du Sénégal for the approval to conduct this study in the Niokolo-Koba National Park. We especially thank the former conservateur of the park, Jacques Gomis, for his support. We are grateful to all the CRP Simenti staff and field assistants for their support in the field.

## Data availability

All data and scripts are available from the corresponding author upon request.

## Ethics statement

The data presented in this study were obtained exclusively through non-invasive camera-trapping.

