## Supplementary material for "Diversity and spatiotemporal activity patterns of medium and large mammals in the Niokolo-Koba National Park, Senegal"

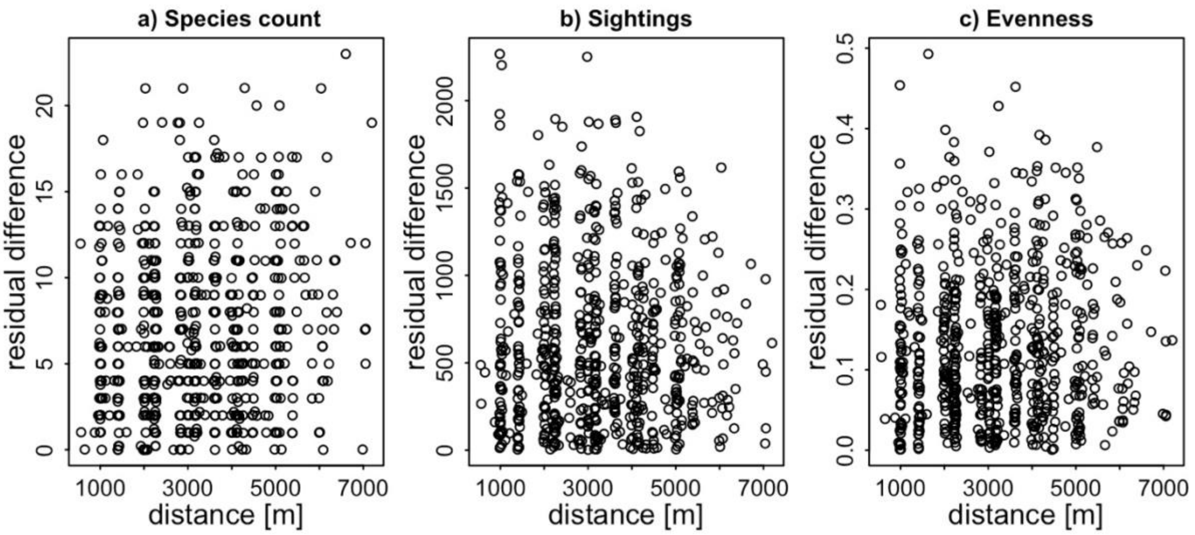


Figure S1: Relationship between pairwise spatial distance and pairwise residual difference for a) species count, b) number of sightings, and c) evenness. Each point represents a unique pair of sampling sites (n = 666). Spatial autocorrelation would result in a clear clustering of residual differences with spatial distance. Pearson correlation coefficients were low (species count: $r = 0.127$; sightings: $r = -0.039$; evenness: $r = 0.042$), suggesting little to no spatial autocorrelation.

In the following Figures S2 to S38, the number of individuals detected per species per sampling site within 100 days of camera trapping across the study site from February 2022 to March 2023 are given.

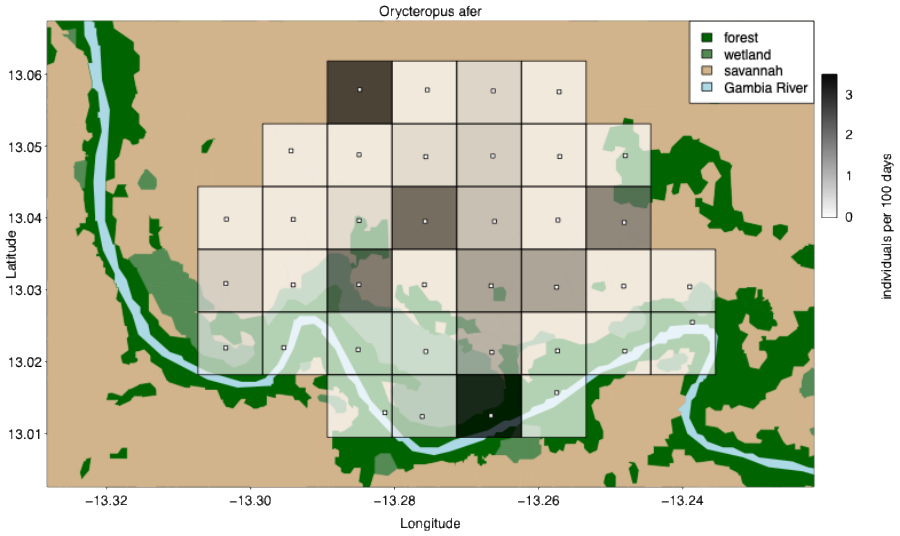


Figure S2**:** Number of individuals of *Orycteropus afer*.

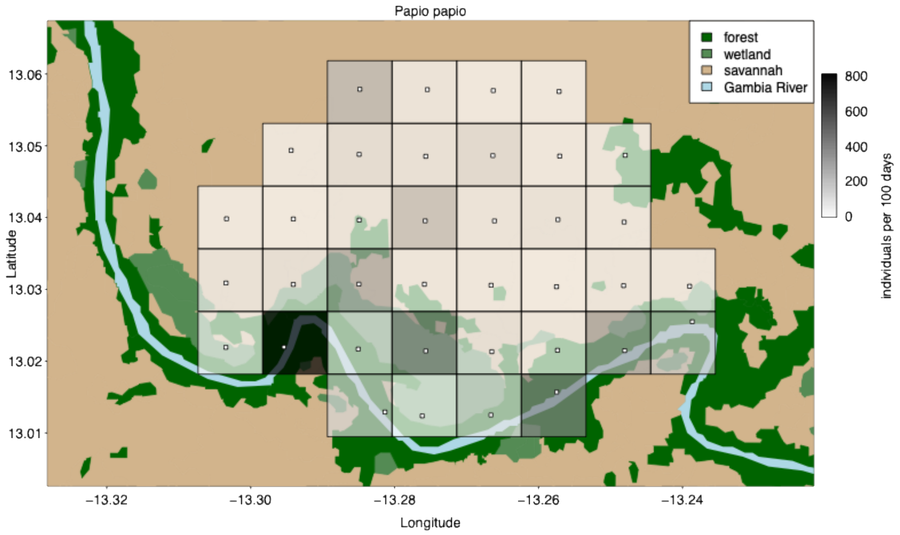


Figure S3: Number of individuals of *Papio papio*.

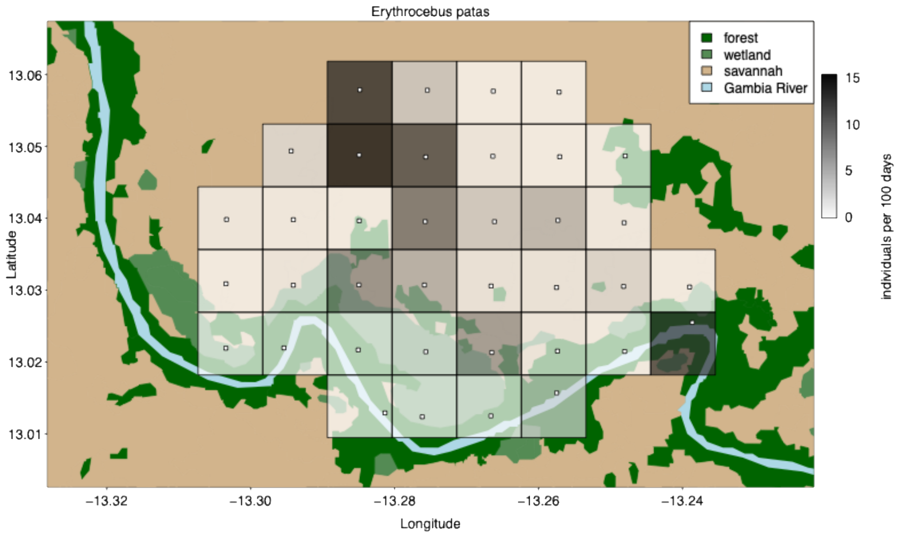


Figure S4: Number of individuals of *Erythrocebus patas*.

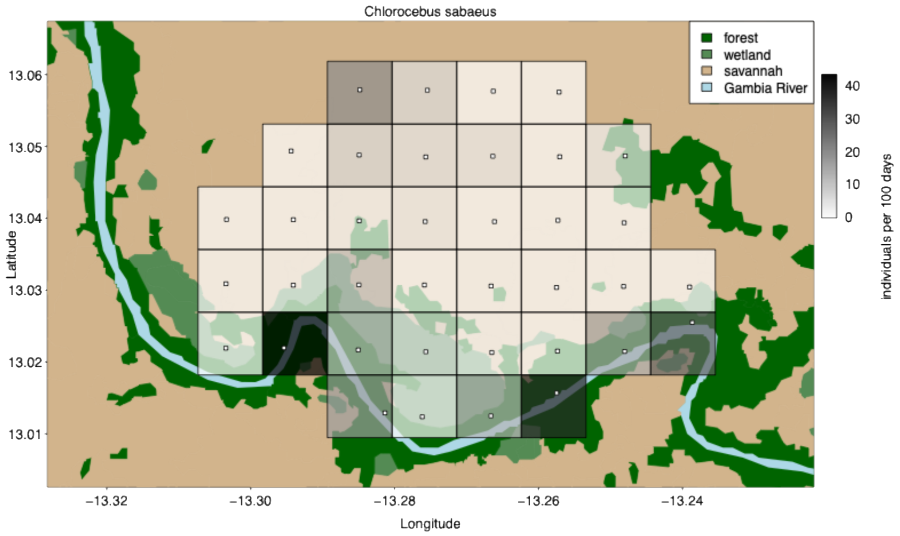


Figure S5: Number of individuals of *Chlorocebus sabaeus*.

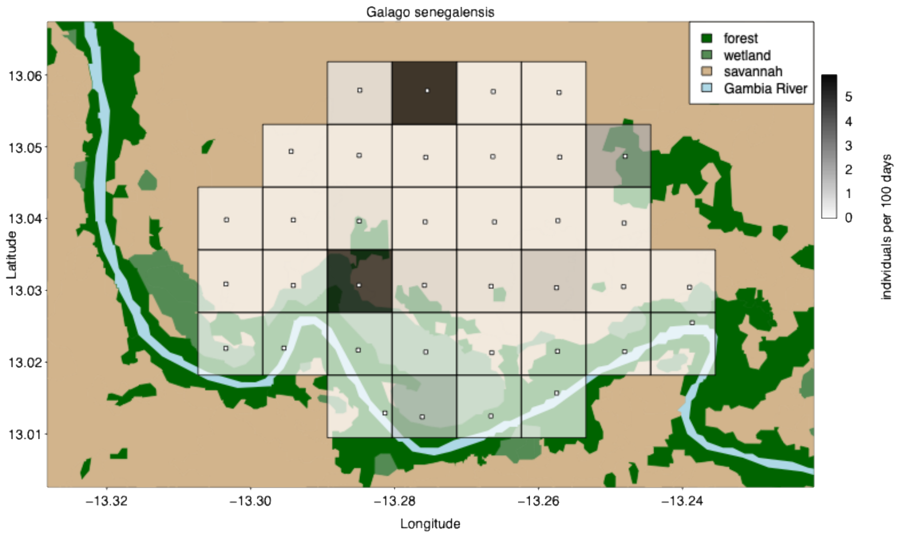


Figure S6: Number of individuals of *Galago senegalensis*.

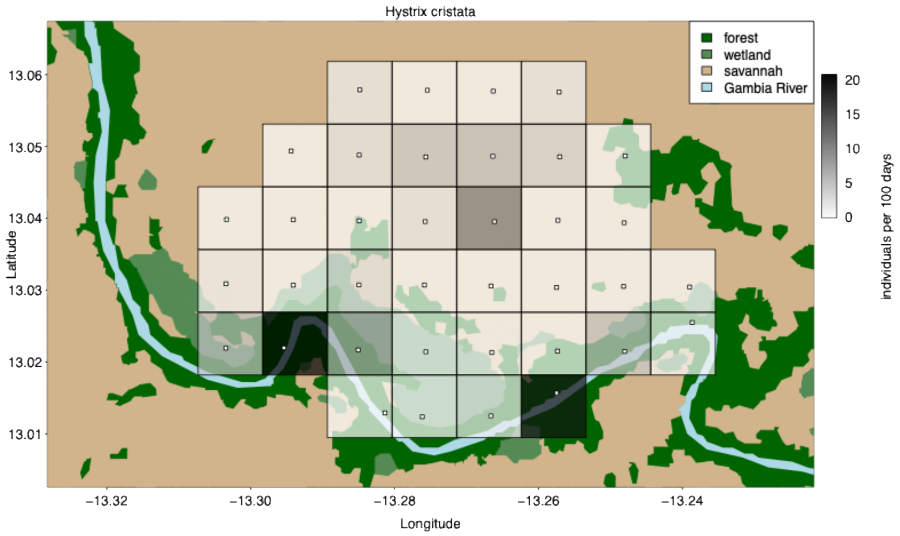


Figure S7: Number of individuals of *Hystrix cristata*.

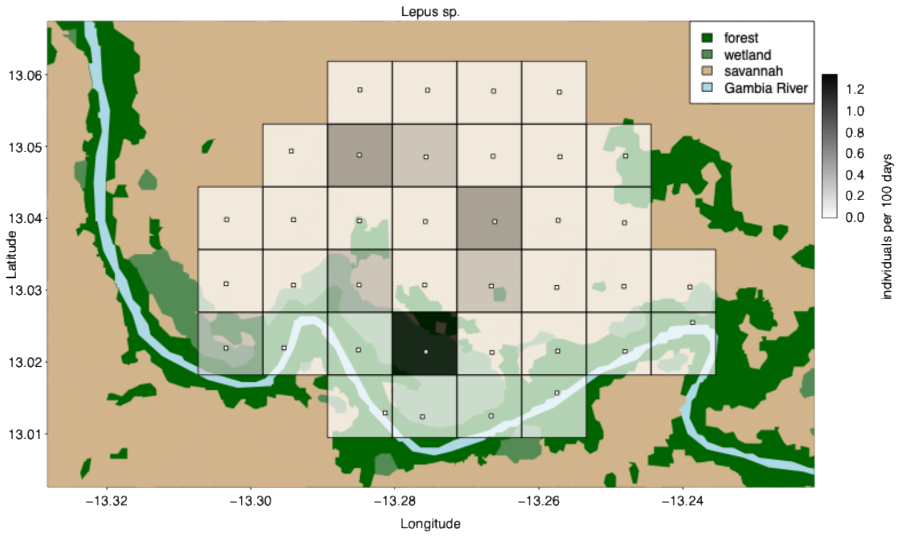


Figure S8: Number of individuals of *Lepus* sp..

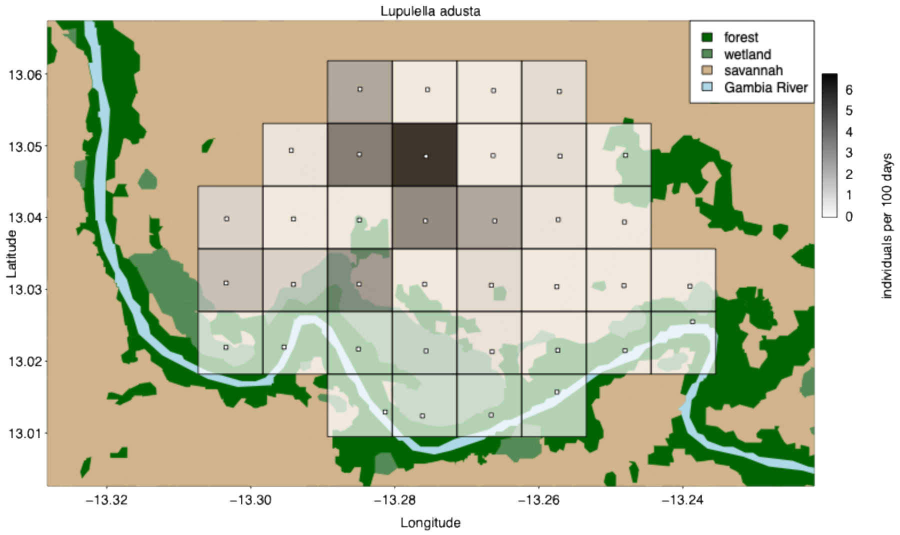


Figure S9: Number of individuals of *Lupulella adusta*.

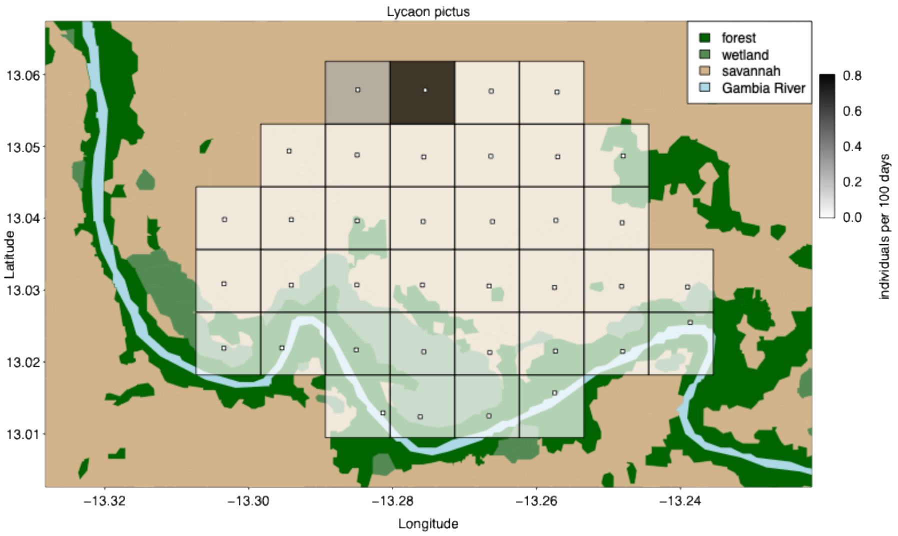


Figure S10: Number of individuals of *Lycaon pictus*.

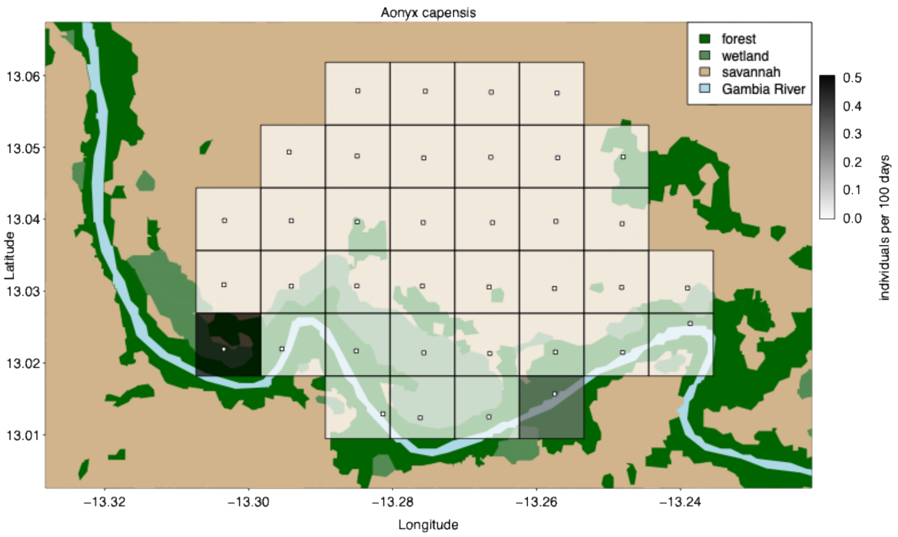


Figure S11: Number of individuals of *Aonyx capensis*.

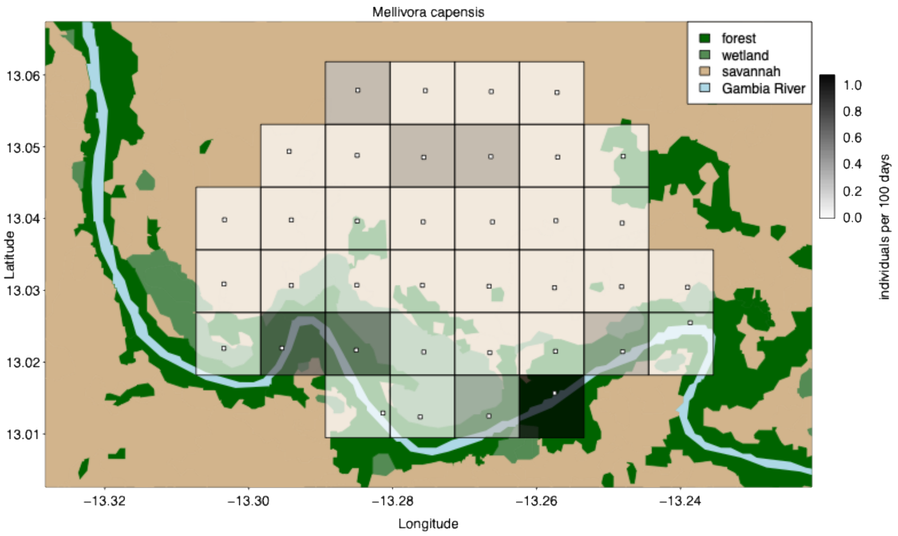


Figure S12: Number of individuals of *Mellivora capensis*.

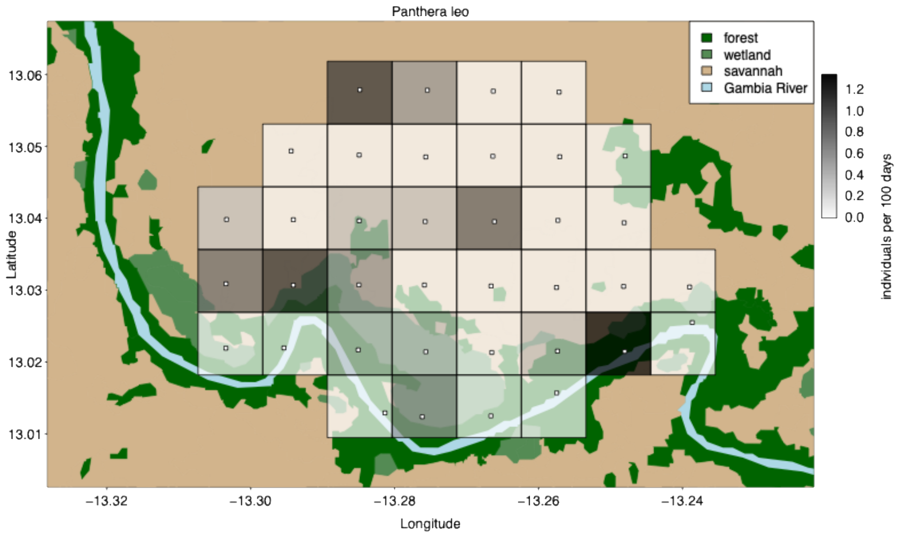


Figure S13: Number of individuals of *Panthera leo*.

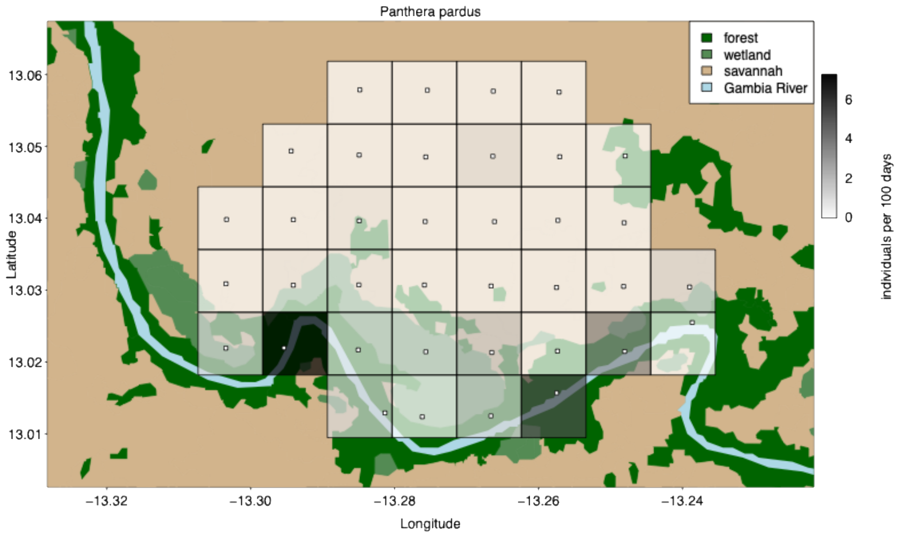


Figure S14: Number of individuals of *Panthera pardus*.

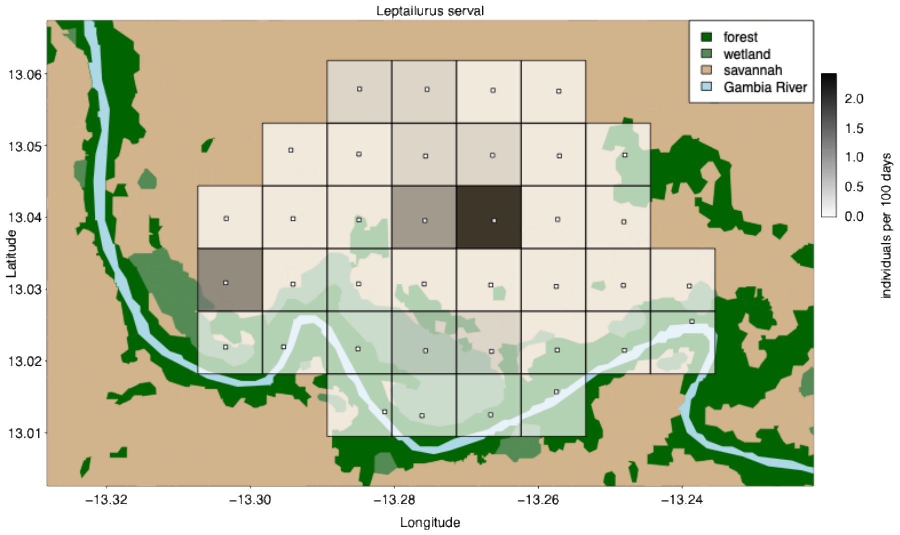


Figure S15: Number of individuals of *Leptailurus serval*.

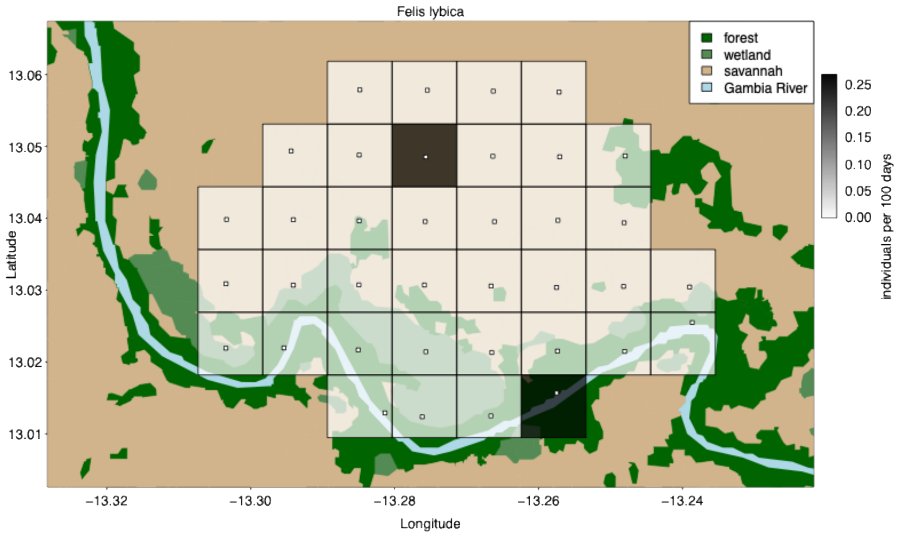


Figure S16: Number of individuals of *Felis lybica*.

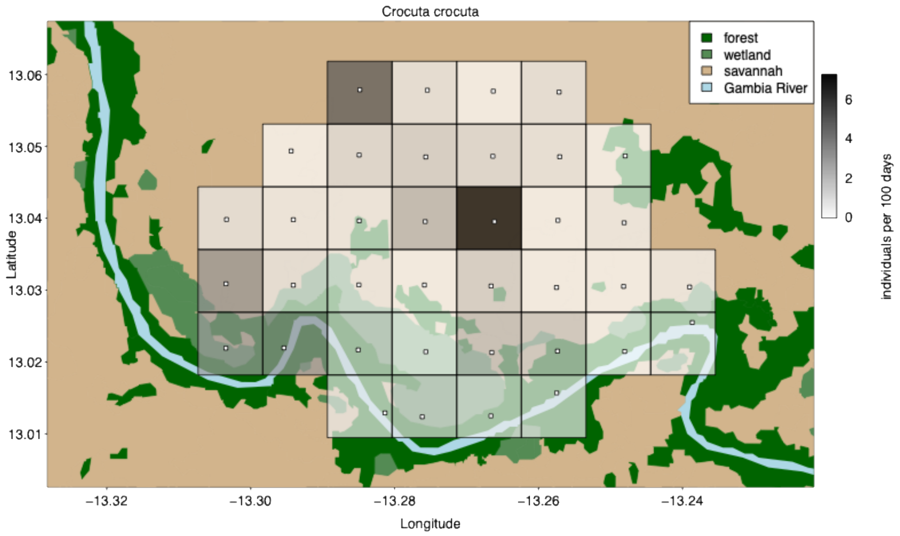


Figure S17: Number of individuals of *Crocuta crocuta*.

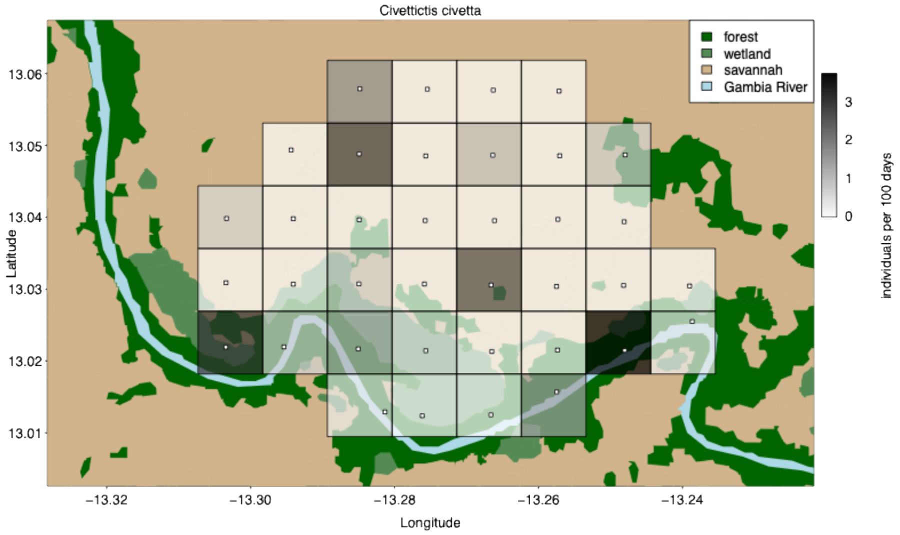


Figure S18: Number of individuals of *Civettictis civetta*.

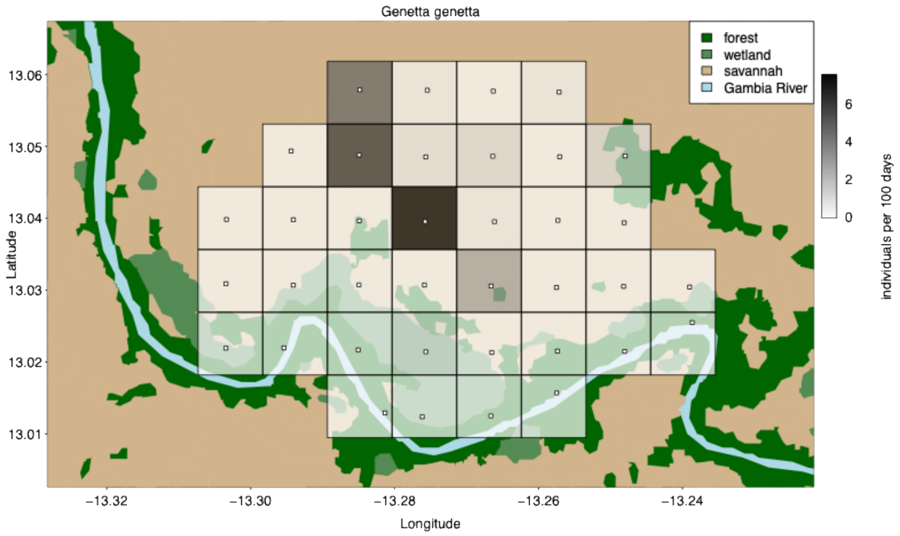


Figure S19: Number of individuals of *Genetta genetta*.

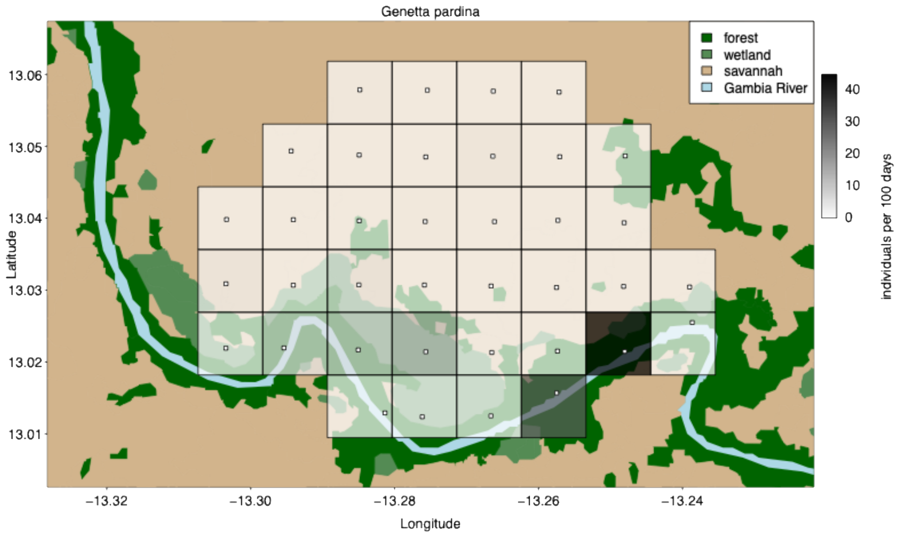


Figure S20: Number of individuals of *Genetta pardina*.

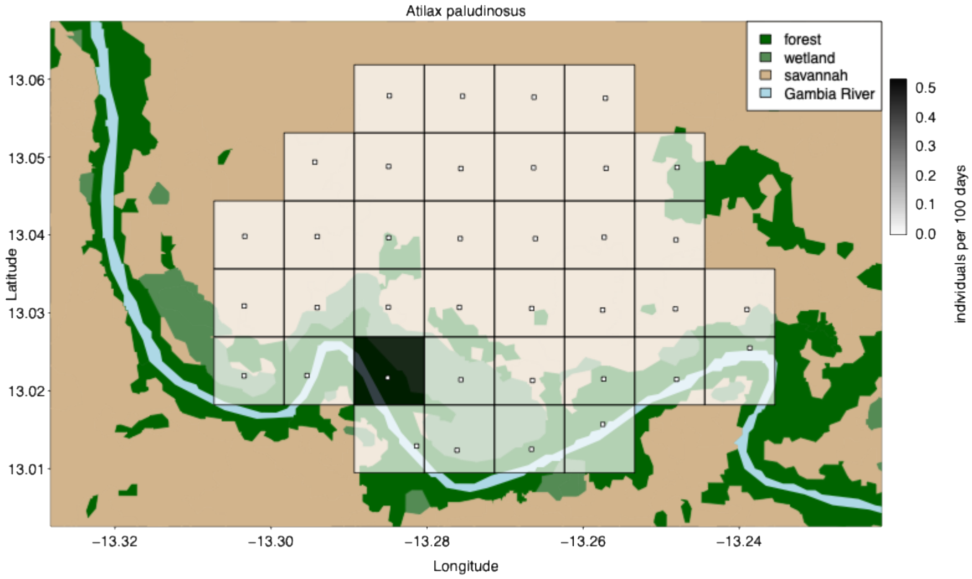


Figure S21: Number of individuals of *Atilax paludinosus*.

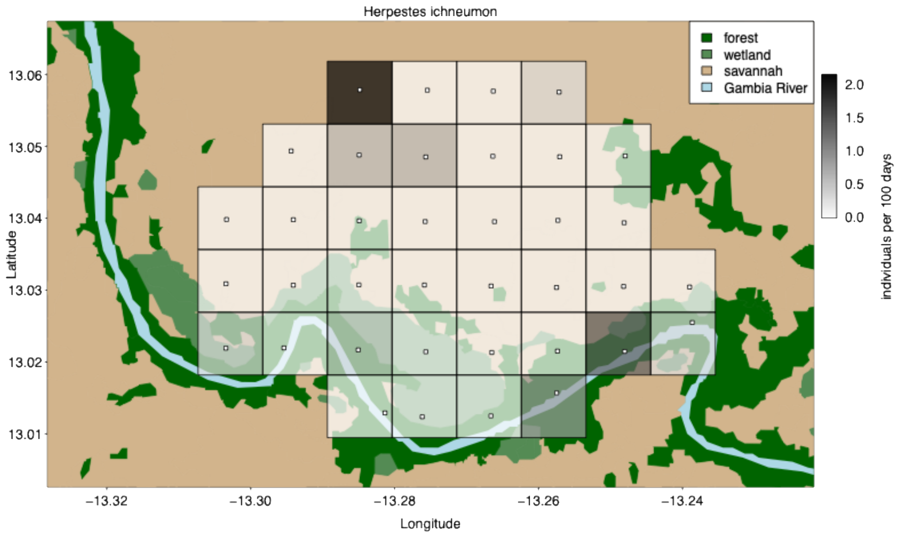


Figure S22: Number of individuals of *Herpestes ichneumon*.

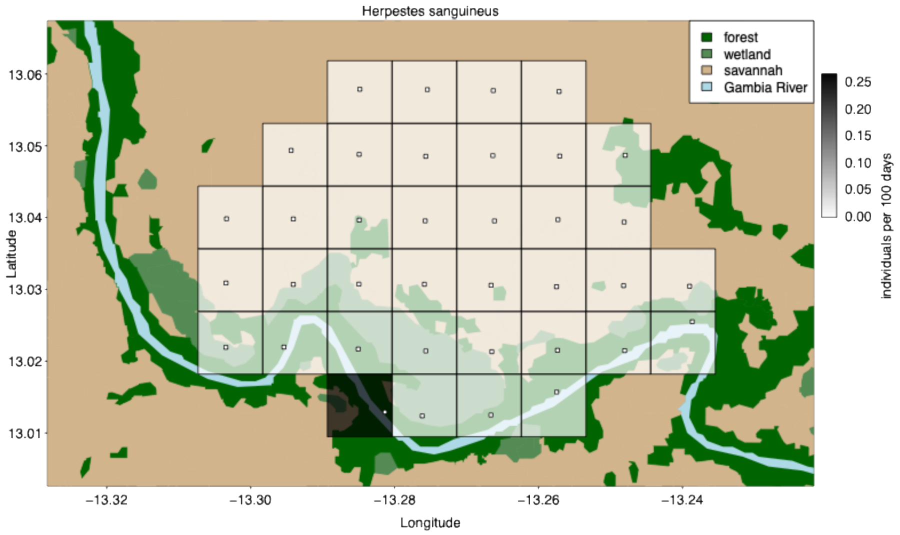


Figure S23: Number of individuals of *Herpestes sanguineus*.

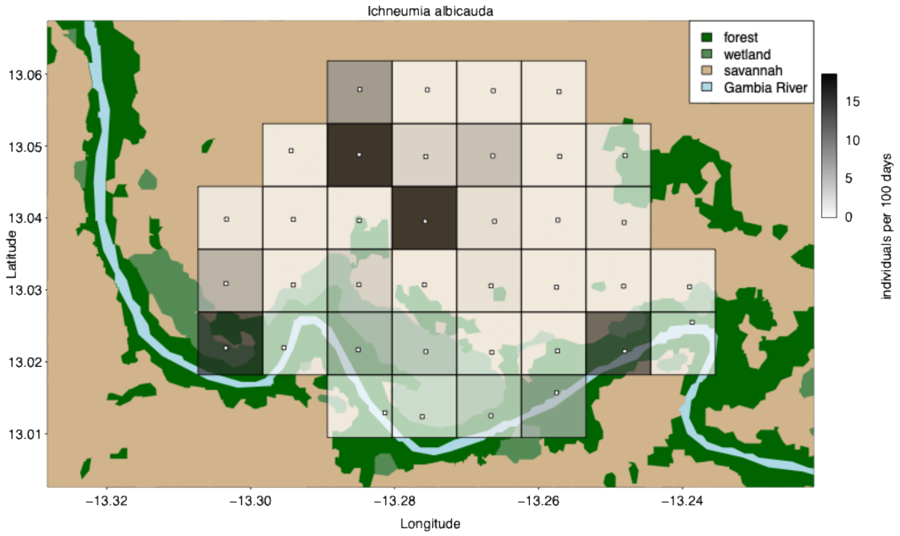


Figure S24: Number of individuals of *Ichneumia albicauda*.

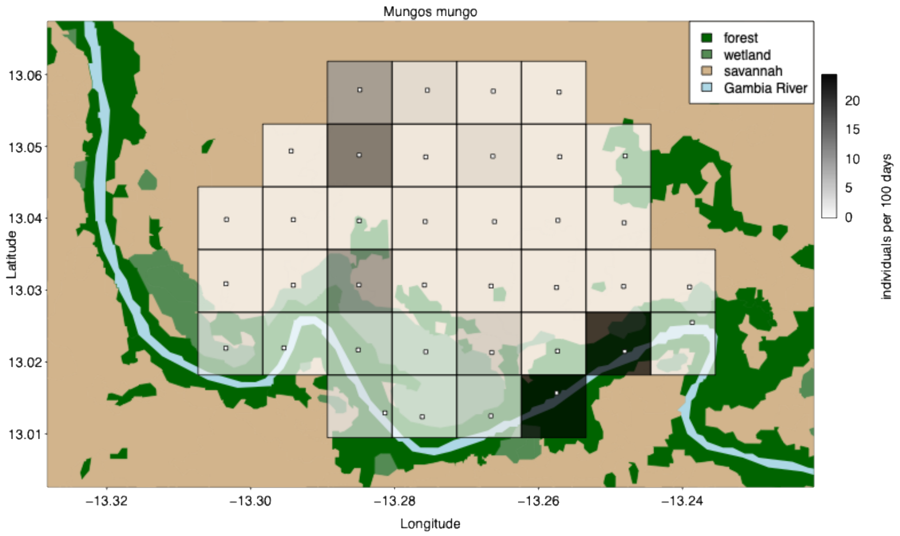

Figure S25: Number of individuals of *Mungos mungo*.

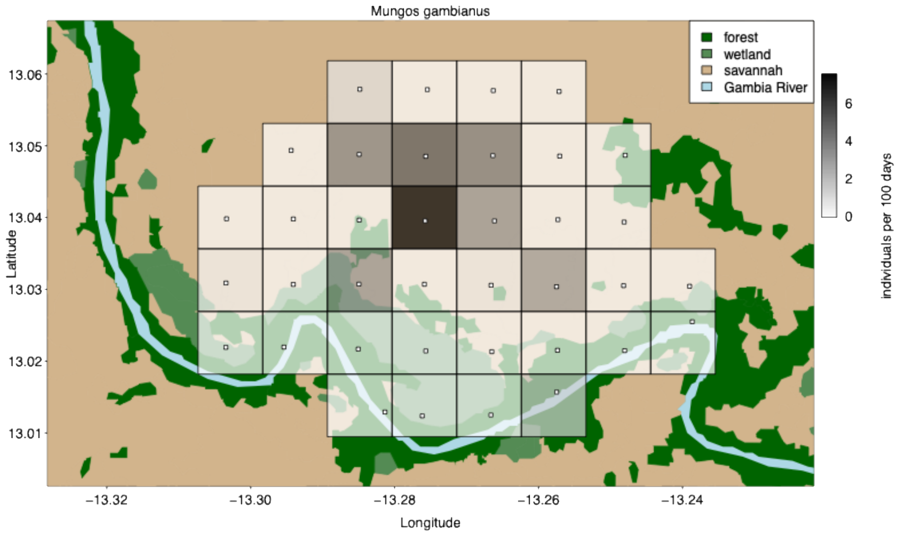

Figure S26: Number of individuals of *Mungos gambianus*.

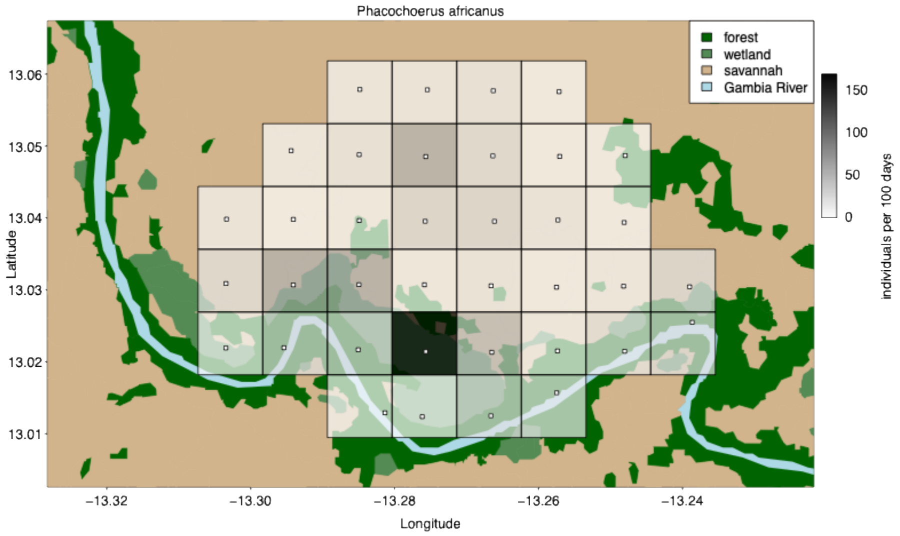

Figure S27: Number of individuals of *Phacochoerus africanus*.

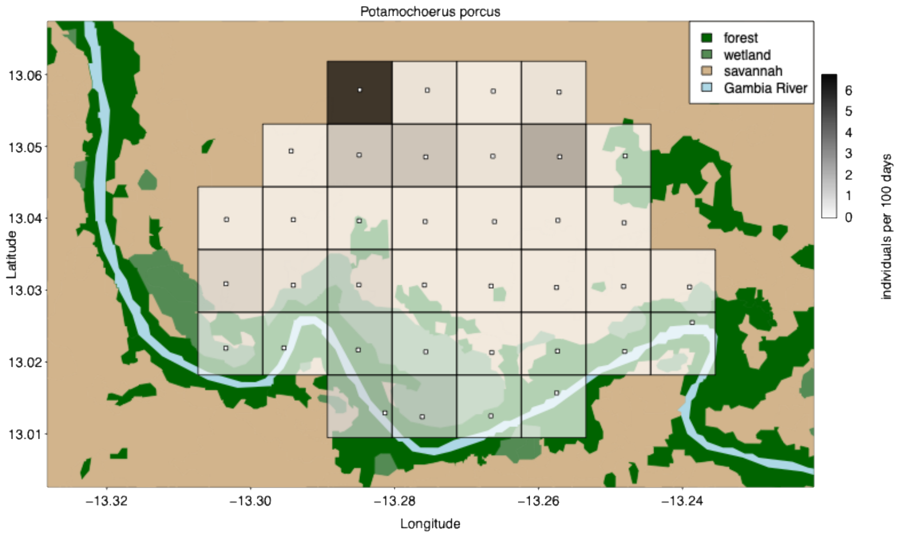


Figure S28: Number of individuals of *Potamochoerus porcus*.

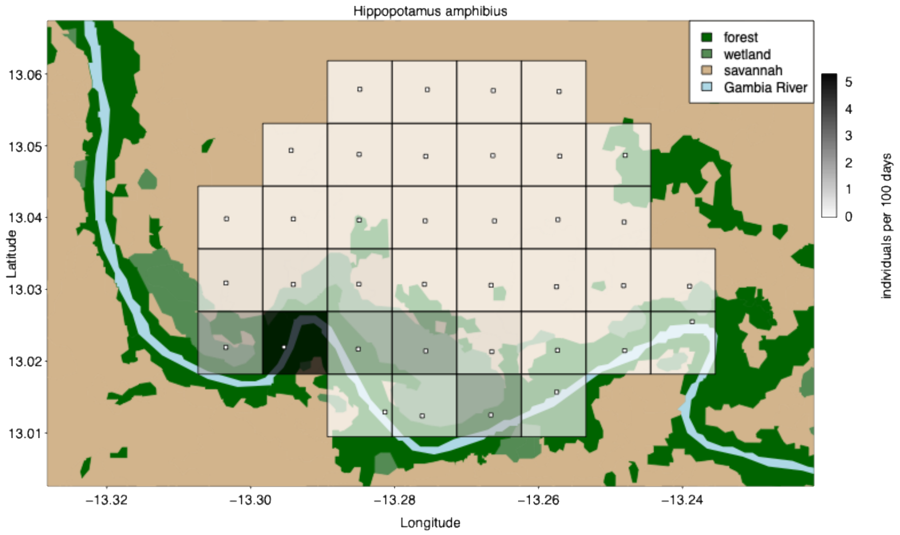


Figure S29: Number of individuals of *Hippopotamus amphibius*.

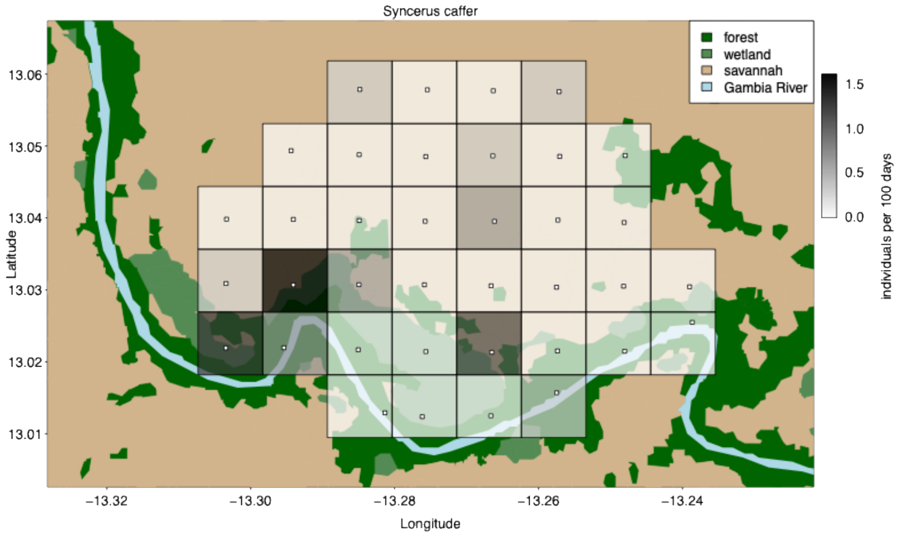


Figure S30: Number of individuals of *Syncerus caffer*.

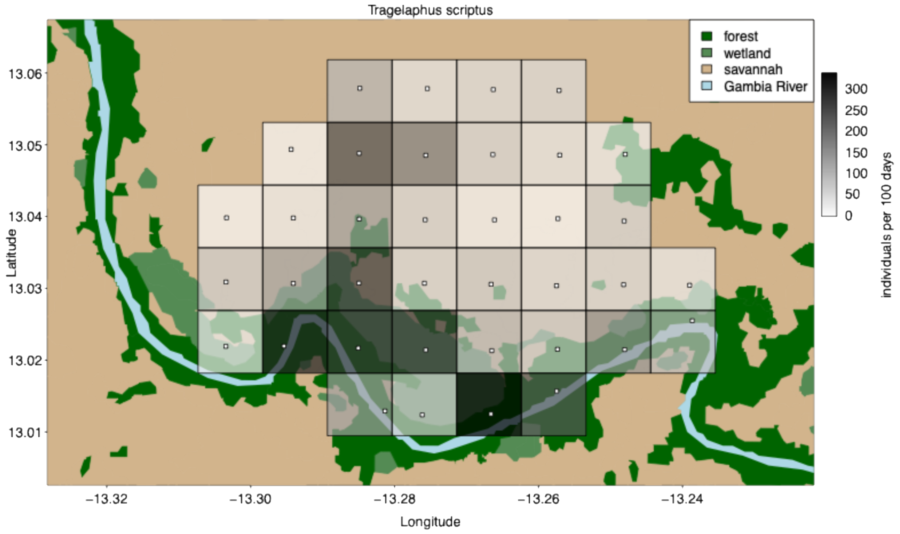


Figure S31: Number of individuals of *Tragelaphus scriptus*.

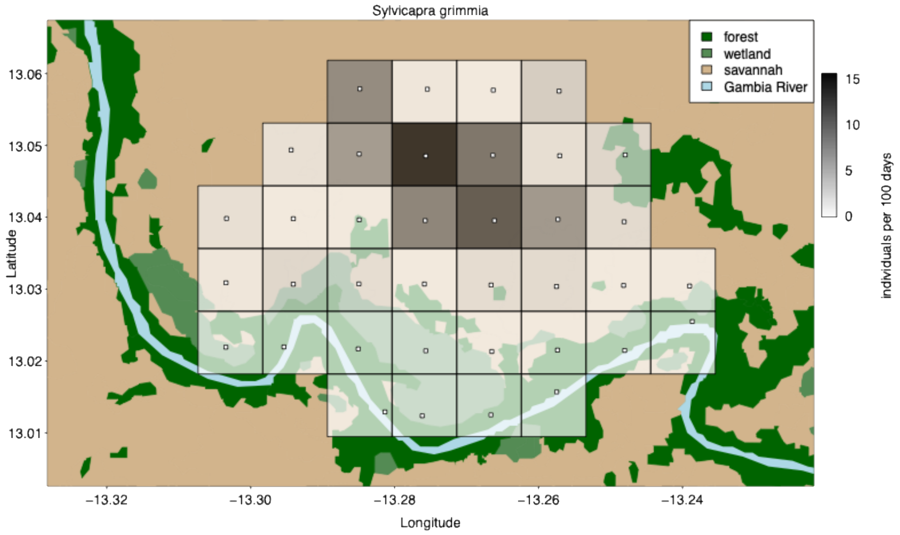


Figure S32: Number of individuals of *Sylvicapra grimmia*.

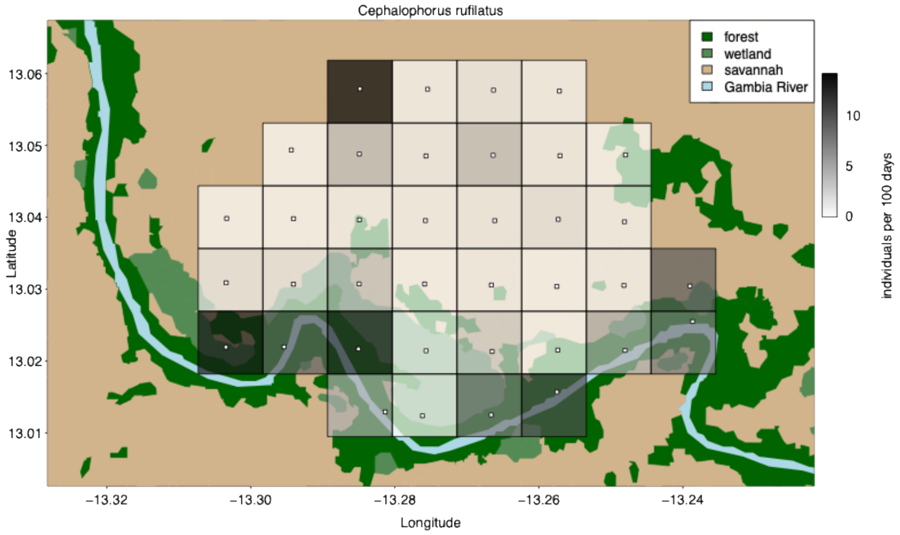


Figure S33: Number of individuals of *Cephalophorus rufilatus*.

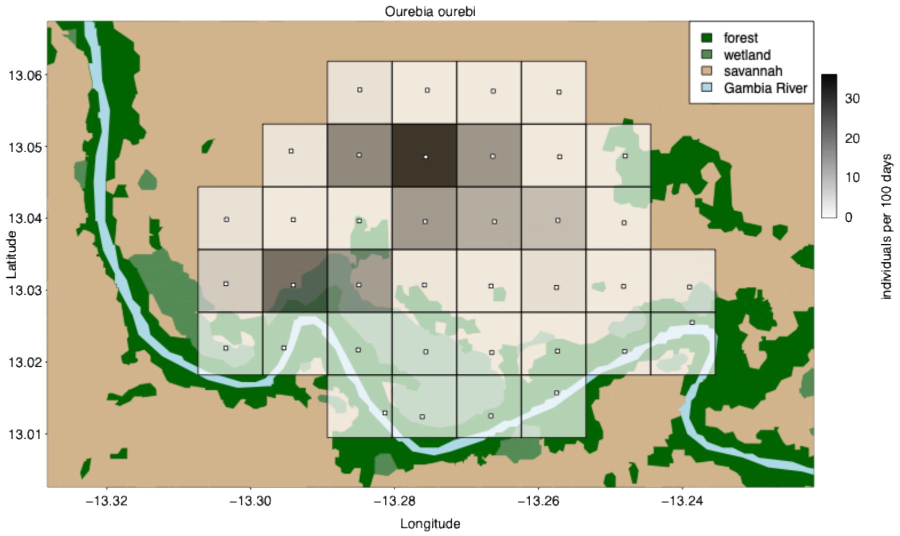


Figure S34: Number of individuals of *Ourebia ourebi*.

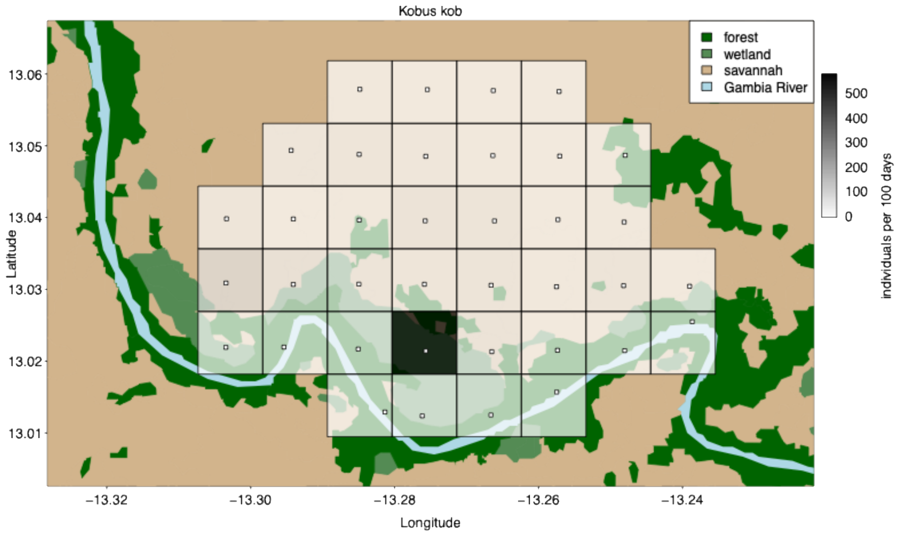


Figure S35: Number of individuals of *Kobus kob kob*.

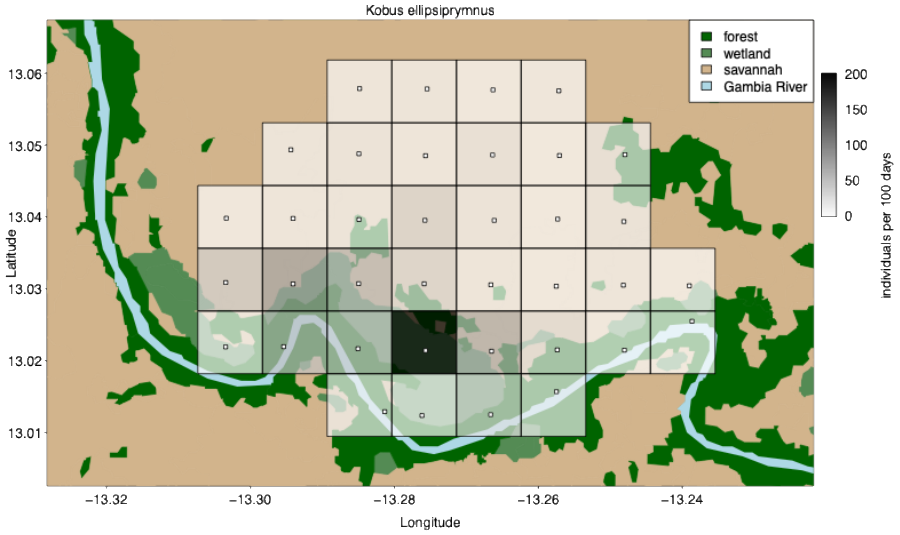


Figure S36: Number of individuals of *Kobus ellipsiprymnus defassa*.

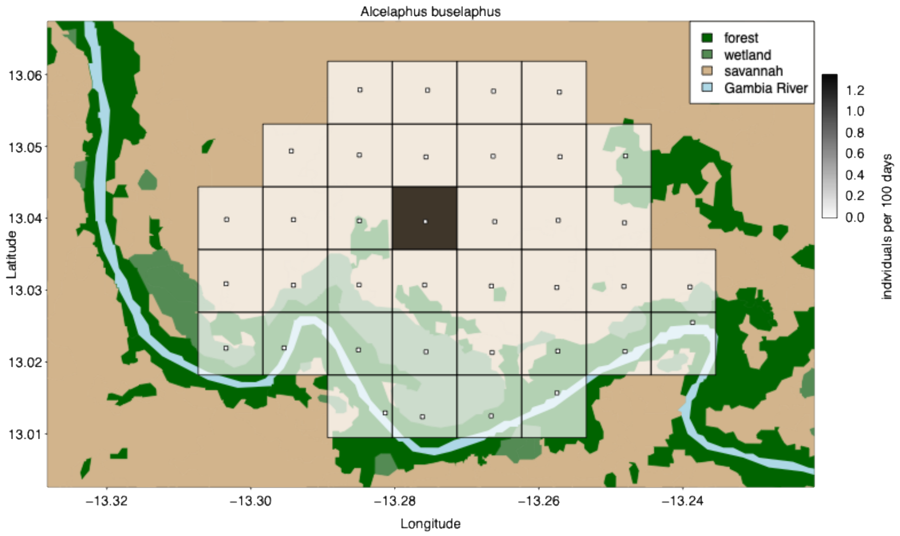


Figure S37: Number of individuals of *Alcelaphus buselaphus major*.

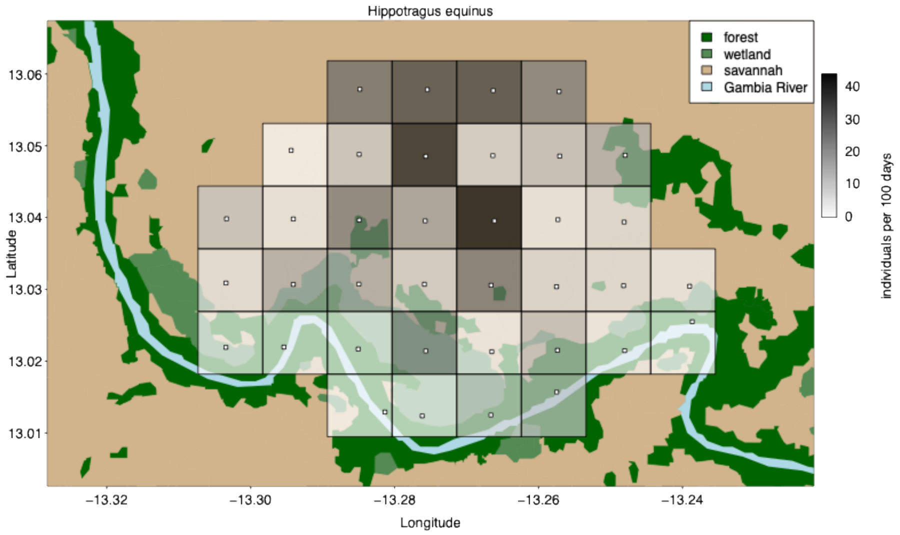


Figure S38: Number of individuals of *Hippotragus equinus koba*.

Table S1: Relative sighting frequencies across habitat types and seasons for species with at least 25 total records from February 2022 to March 2023. Gradual shading of cells reflects the proportion of sightings per habitat and season (white = 0, dark green = 1), providing a visual guide alongside the printed values.

| Species | forest | | savannah | | wetland | |
|---|---|---|---|---|---|---|
| | dry | wet | dry | wet | dry | wet |
| *Orycteropus afer* | 0.11 | 0.27 | 0.89 | 0.73 | 0.00 | 0.00 |
| *Papio papio* | 0.56 | 0.66 | 0.40 | 0.32 | 0.04 | 0.02 |
| *Erythrocebus patas* | 0.15 | 0.03 | 0.83 | 0.97 | 0.02 | 0.00 |
| *Chlorocebus sabaeus* | 0.72 | 0.75 | 0.26 | 0.25 | 0.02 | 0.00 |
| *Galago senegalensis* | 0.06 | 0.13 | 0.94 | 0.88 | 0.00 | 0.00 |
| *Hystrix cristata* | 0.67 | 0.64 | 0.33 | 0.36 | 0.00 | 0.00 |
| *Lupulella adusta* | 0.00 | 0.00 | 0.99 | 1.00 | 0.01 | 0.00 |
| *Panthera leo* | 0.17 | 0.43 | 0.78 | 0.57 | 0.06 | 0.00 |
| *Panthera pardus* | 0.88 | 0.67 | 0.05 | 0.29 | 0.07 | 0.04 |
| *Crocuta crocuta* | 0.40 | 0.13 | 0.60 | 0.86 | 0.00 | 0.02 |
| *Civettictis civetta* | 0.59 | 0.20 | 0.41 | 0.70 | 0.00 | 0.10 |
| *Genetta genetta* | 0.00 | 0.00 | 0.99 | 0.92 | 0.01 | 0.08 |
| *Genetta pardina* | 0.85 | 0.80 | 0.06 | 0.05 | 0.09 | 0.16 |
| *Ichneumia albicauda* | 0.39 | 0.32 | 0.59 | 0.64 | 0.02 | 0.03 |
| *Mungos mungo* | 0.56 | 0.54 | 0.44 | 0.46 | 0.00 | 0.00 |
| *Mungos gambianus* | 0.00 | 0.08 | 1.00 | 0.92 | 0.00 | 0.00 |
| *Phacochoerus africanus* | 0.22 | 0.19 | 0.66 | 0.45 | 0.12 | 0.36 |
| *Potamochoerus porcus* | 0.25 | 0.13 | 0.75 | 0.88 | 0.00 | 0.00 |
| *Hippopotamus amphibius* | 0.79 | 0.71 | 0.07 | 0.29 | 0.14 | 0.00 |
| *Tragelaphus scriptus* | 0.45 | 0.40 | 0.51 | 0.49 | 0.03 | 0.10 |
| *Sylvicapra grimmia* | 0.00 | 0.00 | 1.00 | 1.00 | 0.00 | 0.00 |
| *Cephalophorus rufilatus* | 0.71 | 0.35 | 0.25 | 0.49 | 0.05 | 0.16 |
| *Ourebia ourebi* | 0.01 | 0.00 | 0.99 | 1.00 | 0.01 | 0.00 |
| *Kobus kob* | 0.02 | 0.02 | 0.39 | 0.86 | 0.58 | 0.12 |
| *Kobus ellipsiprymnus* | 0.33 | 0.16 | 0.51 | 0.54 | 0.16 | 0.30 |
| *Hippotragus equinus* | 0.11 | 0.01 | 0.86 | 0.98 | 0.03 | 0.00 |
